# Core and Flanking bHLH-PAS:DNA interactions mediate specificity and drive obesity

**DOI:** 10.1101/2022.02.01.475276

**Authors:** DC Bersten, AE Sullivan, D McDougal, J Breen, RL Fitzsimmons, GEO Muscat, S Pederson, JB Bruning, CM Fan, PQ Thomas, DL Russell, DJ Peet, ML Whitelaw

## Abstract

The basic-Helix-Loop-Helix Per-Arnt-Sim (PAS) homology domain (bHLH-PAS) transcription factor (TF) family comprises critical biological sensors of physiological (hypoxia, tryptophan metabolites, neuronal activity, and appetite) and environmental (diet derived metabolites and environmental pollutants) stimuli to regulate genes involved in signal adaptation and homeostasis^1^. bHLH TFs bind DNA as homo or heterodimers via E-box (CANNTG) response elements, however the DNA binding specificity of the PAS domain-containing bHLH subfamily remains unresolved^1^. We systematically analysed cognate DNA binding hierarchies of prototypical bHLH-PAS family members (ARNT, ARNT2, HIF1α, HIF2α, AhR, NPAS4, SIM1) and demonstrate distinct core (NNCGTG) specificities for different heterodimer classes. The results also show that bHLH-PAS TFs bind over a large footprint 12-15bp and recognise preferential DNA sequences flanking the core. For example, specificity beyond otherwise identical core binding by SIM1 and the HIFs is mediated through N-terminal HIFα-DNA interactions. We also reveal an intimate relationship between DNA shape and both core and flanking TF binding allowing motif sequence flexibility and underpinning TF binding specificity. Furthermore, DNA-shape affinity relationships revealed that novel downstream PAS-A-loop DNA interactions are associated with AT-rich sequences that lead to high-affinity binding, and that loss of this function underpins a monogenic cause of human hyperphagic obesity in a recapitulated SIM1.R171H knock-in mouse model.

Importantly, models of protein-DNA binding accurately predict in vivo occupancy, while response element methylation blocks DNA binding and predicts cell type specific chromatin occupancy. These data provide a definitive and accurate map of bHLH-PAS TF specificity and target selectivity through novel flanking protein-DNA interactions that are crucial for *in vivo* biological function.

## Summary

Gene regulation is mediated by DNA binding transcription factors (TF), via distinct DNA response elements. While consensus DNA sequences play a key role in determining genome occupancy and target gene selection *in vivo,* multiple additional mechanisms must contribute to specificity of TF binding ^2–4^. To investigate TF DNA binding specificity and chromatin selectivity we systematically analysed the mechanisms underlying bHLH-PAS TF family DNA binding.

The bHLH-PAS transcription factor family represents an important model to investigate the mechanisms underlying transcription factor specificity as the family members bind both shared and distinct response elements^1^, display cell-type specific chromatin occupancy and gene expression^5–7^, yet perform distinct biological processes. For example, the hypoxic inducible factors (HIF1a and HIF2a) display non-overlapping biological roles, cell-type or isoform (HIF1a vs HIF2a) specific chromatin occupancy and target gene regulation by unresolved mechanisms^5, 8^. In addition, NPAS4 displays promiscuous response element DNA binding^7, 9^ and performs opposing roles in inhibitory and excitatory neurons via target gene regulation to collectively control neuronal network activity. Lastly, SIM1, SIM2, NPAS1, NPAS3, HIF1a and HIF2a all recognize RACGTG core consensus sequences but perform distinct biological roles in appetite control and hypothalamic development^1^ (SIM1 and SIM2), inhibitory neuron development and activity^10^ (NPAS1 and NPAS3) and the hypoxic response^5^ (HIF1a and HIF2a) indicating that additional encoded specificity is yet to be described.

To explore DNA binding characteristics, bHLH-PAS TF members were profiled by high-throughput DNA binding assays and coupled computational analyses to determine their inherent DNA binding specificities. Through comparative analysis of inherent heterodimer response elements specificity, *in vivo* chromatin occupancy and DNA methylation patterns, DNA shape-affinity relationships, protein-DNA structural analysis, and a novel Sim1^R171H^ knock-in mouse model of obesity we reveal mechanisms underlying TF specificity of the bHLH-PAS TF family.

Initially, *in vitro* DNA binding preferences of several demarcated bHLH-PASA-PASB purified dimers (ARNT/ARNT, ARNT2/ARNT2, AhR/ARNT, HIF1α/ARNT, HIF2α/ARNT, SIM1/ARNT2, and NPAS4/ARNT2) were determined using SELEX sequencing (SELEX-seq)^11, 12^ (Supplementary Fig. 1A-B). The SELEX libraries contained a random 18mer library or libraries containing a fixed E-Box-like (CGTG) core flanked by 8 random nucleotides upstream and 10 nucleotides downstream (Fig. 1A), in an attempt to capture the full specificity upstream and downstream of the core-response element (NNCGTG; core nucleotides 1-6 = c1-c6). By comparing sequence-affinity relationships for all transcription factor complexes using the two approaches of Kmer affinity tables^11^ and No Read Left Behind (NRLB) energy models^13^, high concordance in affinities was found in multiple rounds of selection, and in the relative affinities and TF selectivity from different library strategies (Supplementary Fig. 2A-J).

**Figure 1.**
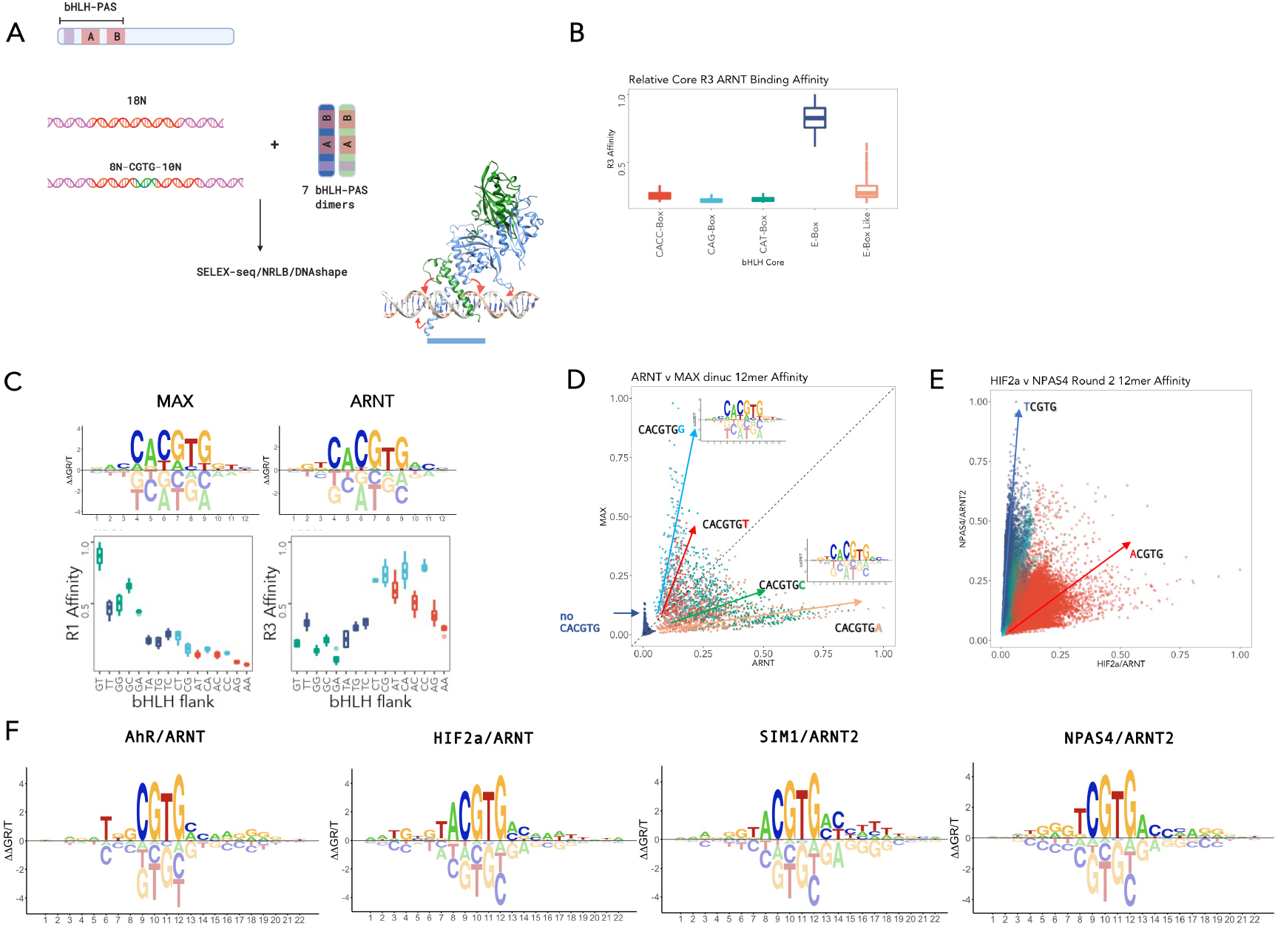
Inherent bHLH-PAS DNA binding affinity and specificity by SELEX-seq. **A)** bHLH-PAS transcription factor dimers were incubated with either random 18mer or FixedCore 8N-CGTG-10N dsDNA ligands prior to selection of bound DNA by Electrophoretic Mobility Shift Assay (EMSA), barcoding and high-throughput sequencing. SELEX high-throughput sequencing was analyzed using a combination of SELEX-seq R, No-Read Left Behind (NRLB) protein-DNA modelling and DNA shape and compared to known and modeled DNA bound bHLH-PAS heterodimeric structures. **B)** Round 3 10mer affinity boxplots (R3 Affinity) of ARNT mediated selection of E-Box (CACGTG) containing probes vs other known bHLH motifs (CACC-Box (CACCTG), CAG-Box (CAGCTG), CAT-Box (CATATG), or E-Box-Like (DNCGTG; D = A, T, G). **C)** MAX vs ARNT derived 12mer energy models (upper panel) reveal E-Box flanking specificity which is also observed in round 3 12mer affinity boxplots (R3 Affinity of palindromic fCACGTGf Kmers) (lower panel) **D)** DNA binding specificity between bHLH subgroups ARNT vs MAX is encoded by E-Box flanking sequences. Affinities of all NNNCANNTGNNN 12mers were scored using dinucleotide NRLB models. Flanking sequences (CANNTG**<u>N</u>**; f_+1_) (or non-CACGTG containing sequences) are colour coded as indicated. **E)** bHLH-PAS transcription factor heterodimer DNA binding can be distinguished by a single nucleotide upstream (N**<u>N</u>**CGTG; c_2_) of the E-Box like core NCGTG. 12mers from SELEX-seq were colour coded for **<u>N</u>**CGTG sequence as indicated. **F)** Energy logos of dinucleotide TF-DNA binding models.

While members of the bHLH superfamily of TFs can bind to various core (CAT-box, CACC-box, CAG-Box, E-box (CACGTG) or E-box-like (DNCGTG)) sequences^2, 14^, we found that ARNT homodimers bound selectively to the E-Box palindrome (CACGTG) while bHLH-PAS heterodimers bound selectively to E-Box like sequences (DNCGTG; D = A,T,G) (Fig. 1B and Supplementary Fig. 2B). Given that E-Box elements are also bound by other non-PAS bHLH TFs such as MAX and MyoD, and that DNA-binding specificity of yeast bHLH TFs can be distinguished by the nucleotides flanking the core E-Box^15^, the DNA binding affinities of MAX vs ARNT were compared. First, protein-DNA binding models from SELEX-seq data were generated and energy logos created representing DNA binding affinities of MAX or ARNT across a 12mer footprint (Fig. 1C). Indeed, comparison of MAX and ARNT flanking specificity of CACGTG containing 12mers or modeled affinities indicated that core flanking nucleotide preferences (ffCANNTGff; f_-1_,f_-2_,f_+1_, f_+2_) were different between ARNT and MAX and suggested selective recruitment of different bHLH TFs factors to distinct extended E-Box elements (Fig. 1C and D).

Within the bHLH-PAS heterodimers selective DNA binding between different complexes was encoded by a single nucleotide at the 5’ of the core (N**N**CGTG or **N**NCGTG; c_1_ or c_2_) (Fig. 1E, Supplementary Fig. 2E-J, and Supplementary 3A-F), except for the HIFα isoforms, where binding affinities were indistinguishable (HIF1α/ARNT vs HIF2α/ARNT; r^2^ = 0.94; Supplementary Fig.2H). Similar to HIFα, poorly selective DNA binding was observed between the closely related homodimers of ARNT vs ARNT2 (r^2^ = 0.78; Supplementary Fig. 2G). We also noted that while AhR/ARNT and NPAS4/ARNT2 exhibit distinct, high affinity core DNA binding motifs, those for SIM1/ARNT2, HIF1α/ARNT and HIF2α/ARNT appear to be indistinguishable (Fig. 1F, Supplementary Fig. 2I-J and Supplementary Fig. 3H-I). However, energy logos (Fig. 1F) indicate that additional specificity encoded in flanking sequences may be sufficient to mediate selectivity between otherwise identical transcription factor binding sites.

To confirm the validity of DNA binding models we used four approaches: 1) comparison of modeled affinities with EMSA gel shifts of flanking variant sequences, 2) qualitative comparison of SELEX derived motifs vs ChIP motif discovery, 3) assessment of the ability of models to predict *in vivo* chromatin occupancy by area under receiver operator curve analysis, and 4) comparison of bHLH-PAS model TF peak scores vs chromatin occupancy strength (Supplementary Fig. 5A-I). Taken together (see supplementary discussion) these analyses supported the validity of DNA binding energy models and SELEX-seq data for prediction of both *in vitro* and *in vivo* DNA binding.

Comparing energy logos from all HIFα/ARNT and SIM1/ARNT2 SELEX-seq experiments demonstrated that different upstream nucleotide preferences exist between HIFα/ARNT and SIM1/ARNT2, with TG nucleotide preference for HIFα/ARNT at f_-3_-f_-4_ upstream of the core sequence (**<u>NN</u>**NNTACGTG) (Fig. 2A). In addition, comparison of HIF2α/ARNT and SIM1/ARNT2 fixed core energy logos indicated that both upstream and downstream nucleotides likely contribute to specificity (Supplementary Fig. 6A-E). This was supported by comparison of kmer affinities of preferential upstream and downstream sequences, demonstrating HIF2α and SIM1 preferential binding to identical core (TACGTG) response elements *in vitro (*Fig. 2B, Supplementary Fig. 6B-E). The *in vitro* specificity of HIFα/ARNT heterodimers was also reflected by increased occupancy of HIFα at GNNTACGTG containing HIF1α ChIP peaks (and less pronounced for HIF2α) in both HepG2^5^ and MCF7 cells^6^, supporting *in vitro* DNA binding preference being conferred to *in vivo* chromatin site selection (Fig. 2C). In addition, linear regression coefficients of HIFα model-based scoring of HIFα ChIP-peaks were higher than SIM1 models and larger than other bHLH-PAS models, indicating that the more intense HIFα/ARNT ChIP peaks are encoded by flanking DNA specificity *in vivo* (Fig. 2D). Taken together, these observations indicate that HIFα/ARNT flanking nucleotide specificity can direct HIF*α in vivo* DNA occupancy, and provides evidence for how target gene specificity between closely related members of the bHLH-PAS family is achieved.

**Figure 2.**
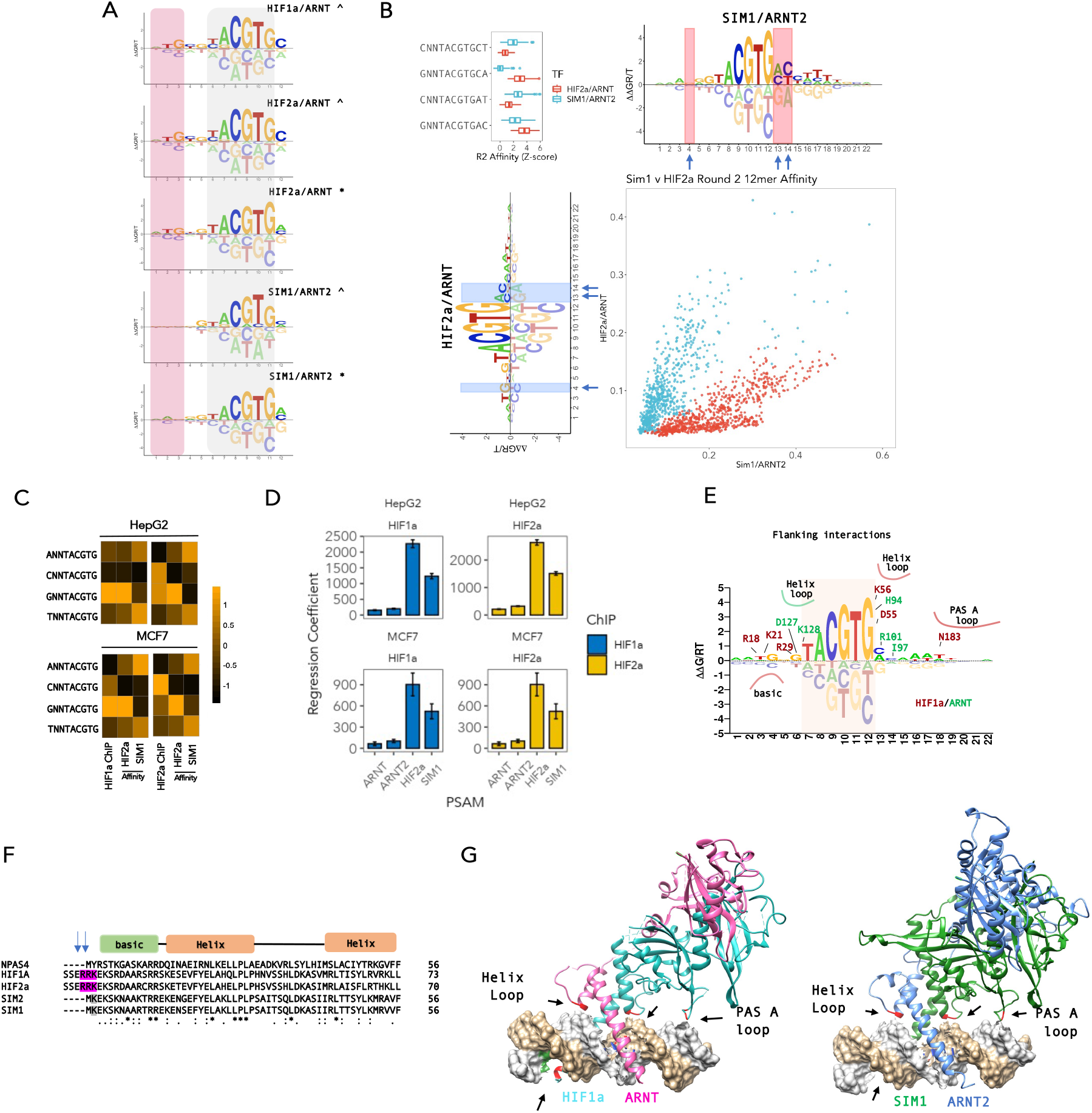
bHLH-PAS DNA binding specificity is encoded by nucleotides flanking the NNCGTG core. **A.)** Upstream nucleotide specificity of SIM1 vs HIFα (HIF1α or HIF2α) from SELEX derived DNA binding energy models of HIFα/ARNT vs SIM1/ARNT2 from either random 18mer (^) or FixedCore 18/22mer (*) SELEX strategies showed that HIFα subunits prefer T and G at position f_-3_ and f_-4_ (upstream of core NNCGTG) vs C and C at position f_-3_ and f_-4_. SIM1 SELEX did not appear to have a strong preference or aversion to nucleotides at position f-3 and f-4. **B.)** Upstream f_-3_ (nucleotide −3 from NNCGTG) and downstream f_+1_-f_+2_ (nucleotide +1 and +2 from NNCGTG) encode specificity between share core binding (TACGTG) SIM1/ARNT2 and HIF2α/ARNT transcription factors. Scatterplot comparing Kmer Affinities (a subset of 12mers) for HIF2α/ARNT vs SIM1/ARNT2 coloured by blue = **<u>G</u>**NNNACGTG or red = **<u>C</u>**NNNACGTG demonstrating preferential DNA selection by SIM1/ARNT2 or HIF2α/ARNT. (Top left) Boxplots comparing selective round 2 12mer kmer affinities (Z-score normalised). Blue arrows indicate most variant positions in energy logos between HIF2a/ARNT and SIM1/ARNT2 **C.)** Upstream f_-3_ HIFα preferential sequence specificity is found at HIFα ChIP peaks in HepG2 (Upper panel) and MCF7 cells (Lower panel). ChIP DNA regions were scored by NRLB energy models for SIM1/ARNT2 and HIF2a/ARNT and mean scores were compared to ChIP peak scores at HIF1α/HIF2α peaks and represented by heatmap. **D.)** Linear regression coefficients (± SE) comparing models of *in vitro* DNA binding to *in vivo* ChIP-seq data. HIF1a (blue) or HIF2a (yellow) ChIP-peak DNA (HepG2 or MCF7) was scored using NRLB Models for HIF2a/ARNT, SIM1/ARNT2, ARNT or ARNT2. **E.)** Extensive protein-DNA contacts contribute to Core (NNCGTG shaded orange) flanking nucleotide specificity. Overlay of HIF2a/ARNT energy model with HIF2a (red) or ARNT (green) amino acid-DNA (Phosphate, sugar, or base) contacts curated from (PDB 4ZPK). Basic, helix or PAS A loop domain interactions are outlined. **F.)** basic amino acids in an N-terminal extension of HIF1α and HIF2α not observed in SIM1 or SIM2 interact with upstream (f_-3_ and f_-4_ from NNCGTG) sites. Blue arrows indicate positions in HIFα that contact the T and G at position −3 and −4 (upstream of core NNCGTG) **G.)** Structures of HIF1α/ARNT and SIM1/ARNT2 on DNA were used to illustrate protein DNA interactions that contribute to flanking nucleotide specificity. Left panel -HIF1a (cyan) ARNT (pink) DNA structure and right panel - SIM1/ARNT2/HRE (AGGC**<u>TG</u>**CG**TACGTG**CGGGT**<u>C</u>**GT; flanking nucleotide contacts underlined) modeled structure (on PBD:4ZPK). Core flanking protein-DNA interactions are outlined with black arrows and red amino acids, upstream HIF1α R18 interaction with T f_-4_ and G f_-3_ (from NNCGTG) (green).

Next, to further investigate the mechanism that directs specificity differences between HIFα/ARNT heterodimers vs SIM1/ARNT2 heterodimers the structures of HIF1α/ARNT/HRE^16^ (PDB: 4ZPK) and modeled structure of SIM1/ARNT2/HRE were compared (Fig. 2E and G). Analysis of protein-DNA interactions revealed extensive contacts upstream and downstream of the core DNA binding sequence (Fig. 2F). This included HIFα Arg18 DNA contacts upstream (f_-3_-f_-4_ of the core binding site, **NN**NNTACGTG) at the optimal site of HIFα DNA binding which result from a N-terminal extension of the basic DNA binding domain that is absent in SIM1 (Fig. 2F and G). This indicates that HIFα may have evolved unique specificity to overcome DNA site competition with other bHLH-PAS TF complexes that preferentially bind a TACGTG core.

While no DNA contacts were identified that could explain the proximal (f_+1_/f_+2_ TACGTG**NN**) downstream specificity between HIFα/ARNT and SIM1/ARNT2, the distal (f_+5_-f_+8_) downstream nucleotide preferences revealed a preference for AT-rich sequences close to PAS loop contacts (Fig. 2G). We hypothesized that distal AT-rich specificity may be indicative of more complex shape requirements to accommodate high-affinity DNA binding. To examine this, kmer affinity tables and energy models were used to investigate nucleotide codependences and DNA shape parameters such as Minor Groove Width (MGW), Propeller Twist (ProT), Helical twist (HelT), or Roll^17^.

High affinity AT-rich downstream sequences (f_+5_-f_+8_) were associated with decreased ProT, and MGW associated with high-affinity binding in both SIM1/ARNT2 and HIF2α/ARNT (Fig. 3A-B and Fig. 4I-J). Structural loop remodeling (see supplementary discussion) enabled PAS A loop extension more distally, allowing Arg/Lys residues in the loop to reorient to come in close proximity or interact with DNA at a conserved position for HIF2α and SIM1 (Fig. 3C-E). Specifically, Arg 181 in HIF2α and Arg 171 in Sim1 extend into the major groove at the site of AT-enrichment (Fig. 3D-E). AT-rich regions are often bound by chromatin associated proteins containing AT-hook domains^18, 19^, Intriguingly, alignment of HIF1α and HIF2α PAS A loop (pal) with AT-Hook motifs from MeCP2 and HMGB1 revealed a conserved RGR AT-hook interaction motif (Supplementary Fig. 7A-B). Mutations in the AT-hook domain of MeCP2 have been shown to lead to Rhett syndrome intellectual disability in humans and mouse models through reduced nucleosome DNA interaction^20–22^, indicating that mutations in other AT-Hook domain containing proteins may be important for disease etiology.

**Figure 3.**
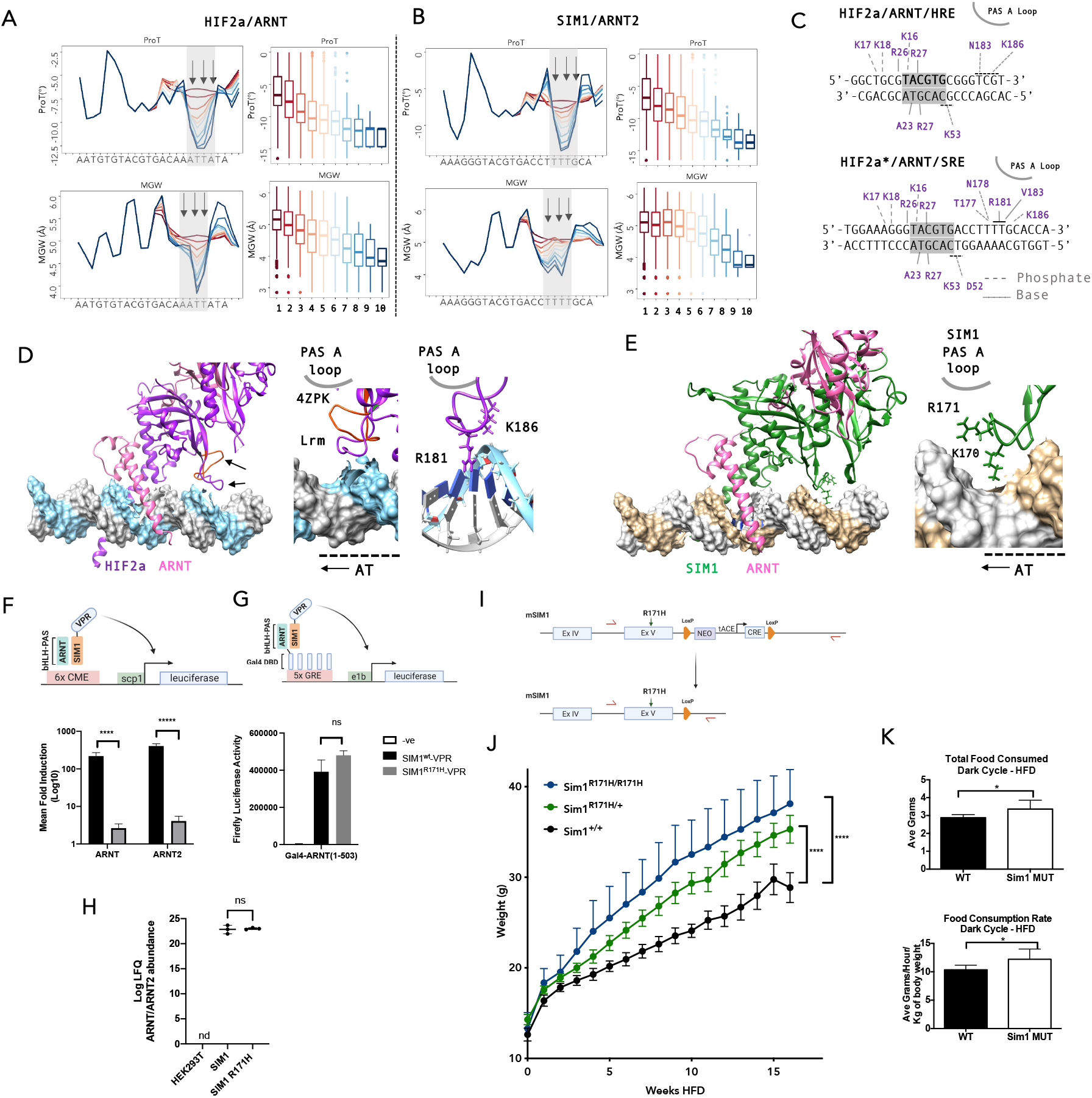
PAS interactions distal to the core binding sites are associated with AT rich, narrowed MGW and reduced ProT revealing SIM1.R171H as a driver of monogenic hyperphagic obesity. **A.)** HIF2a/ARNT or **B.)** SIM1/ARNT2 DNA Shape analysis of mean ProT (top left panel) and MGW (bottom left panel) affinity binned (low = red, blue = high, bins = 0-10) by dinucleotide NRLB model scored downstream 22mers. ProT (top right panel) and MGW (bottom right panel) boxplots at position at T17 (middle arrow; f_+7_). **C.)** HIF2a DNA contacts from PDB 4ZPK or PAS loop remodeled and SRE DNA (28mer) were mapped onto the HRE DNA sequence used in crystallography (DNAProDB), the PAS loop position is indicated for comparison with **D)**, base and phosphate contacts are indicated with solid and dotted lines, respectively. **D.)** HIF2a/ARNT/SRE structure (HIF2a – purple, ARNT – pink) PAS loop remodeling vs crystal structure (4ZPK – red) shows the PAS loop rearrangement to more distally penetrate into the major groove at T16 and T17 (f_+6_ and f _+7_) (correlating with the positions of shape distortion arrows in A.), and AT rich regions identified by NRLB models) allowing R181 and K186 interaction with DNA. **E.)** SIM1/ARNT/SRE structure (HIF2a – purple, ARNT – pink) PAS loop remodeling shows the PAS loop extension into the major groove at T16 and T17 (correlating with the positions of shape distortion arrows in B.) and AT rich regions identified by NRLB models) allowing R171 and K170 interaction with DNA. **F.) and G.)** SIM1(1-348)-VPR or SIM1.R171H(1-348)-VPR were coexpressed with **F.)** ARNT(1-503) or ARNT2(1-455) and a 6xTACGTG luciferase reporter, **G.)** Gal4-ARNT(1-503) and a 5x GRE reporter to assess transcription factor activity and dimerisation, respectively. **F.)** Mean fold induction /ARNT or ARNT2 alone was calculated for each replicate and then the log10 transformed prior to statistical tests. **F.)** and **G.)** Stats were calculated comparing the multiple unpaired t-tests. **** p =0.000626, *****p = 0.000013. n = 3 (± SEM) independent experiments **H.)** Immunopurification (mock (HEK293T) vs SIM1 WT vs SIM.R171H) proteomics (see methods and Supp for details) and label free quantification was used to assess the ability of SIM1 WT or SIM1.R171H to dimerise with ARNT or ARNT2. mean (± 95% CI). nd = not detectable, ns = not significant p = 0.7661 unpaired two tailed t-test. **I.)** SIM1.R171H knock-in mouse model construct and generation. Primer sites are indicated with red arrows. **J.)** mean (±SEM (+ for SIM^R171H/R171H^ for clarity) weight gain of SIM1 WT (n= 8), vs SIM^R171H/+^ (n = 12) vs SIM^R171H/R171H^ (n = 5) littermates over a period of 16 weeks on a high fat diet (HFD). Sim1^+/+^ vs. Sim1^R171H/+^ p = 0.0000007568, WT v Sim1^+/+^ vs. Sim1^R171H/R171H^ p = 0.0000002663, ****p <0.0001. **K.)** (left panel) Average of the total HFD food consumed per mouse during 12 hr period (Dark). (right panel) - Hourly food consumption rate for each mouse, averaged over the 12 hour dark (active) period and corrected for bodyweight. SIM1 WT vs SIM1 Mut (SIM1^R171H/+^) showed increased average food consumption (**2.879 ± 0.06613 vs 3.359 ± 0.1877, n=7 *p=0.0328)*** and increased food consumption rate (**10.34 ± 0.3113 vs 12.21 ± 0.6810, n=7 *p=0.0281*.**) Data analysed by unpaired two-tailed t test.

**Figure 4.**
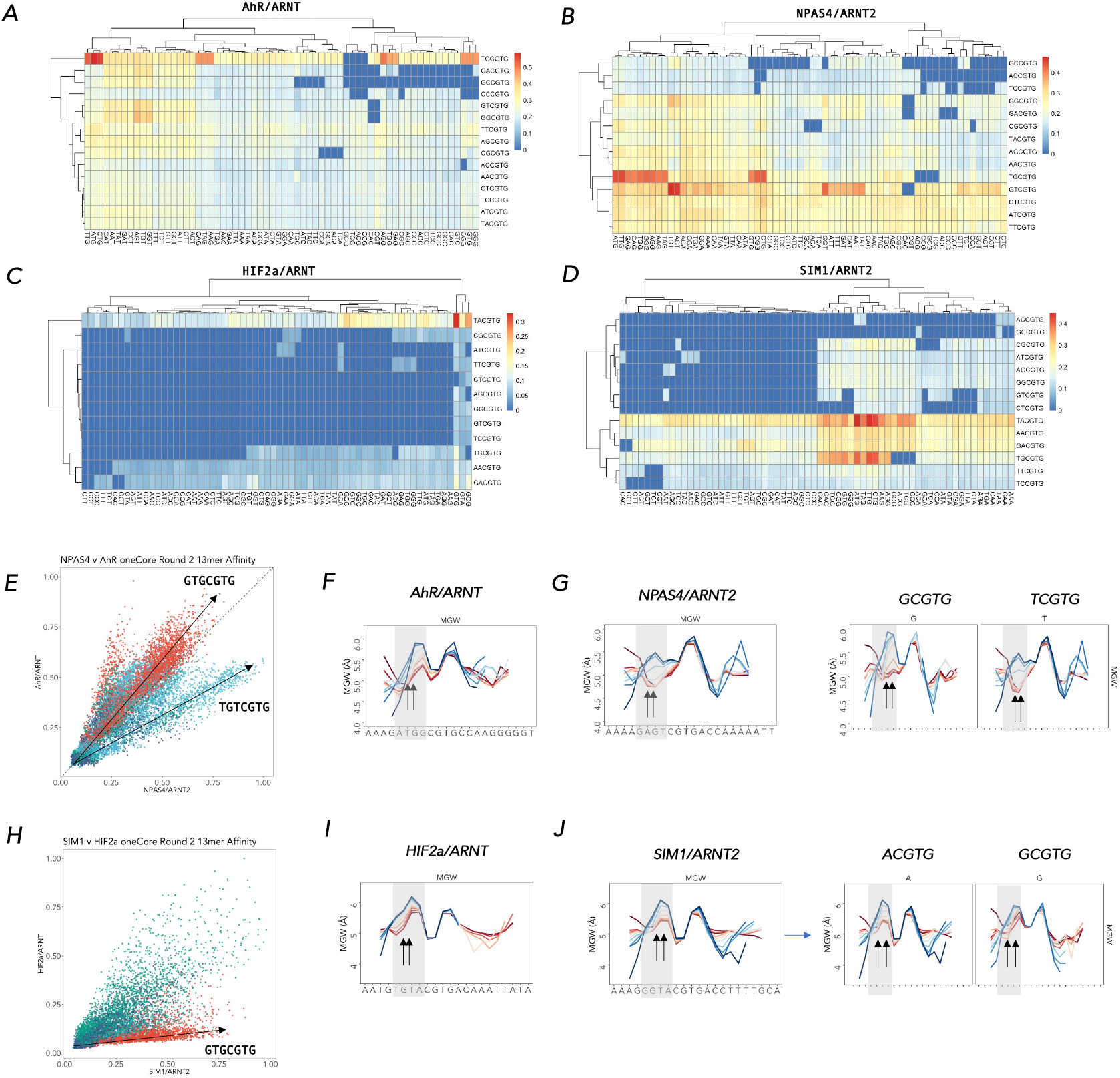
Upstream nucleotide co-dependencies encoded by DNA Shape confers transcription factor selectivity. **A-D.)** 12mer Kmer Affinities (oneCGTG per line) comparing Core NNCGTG (y) and upstream trinucleotides NNNxxCGTG (x). Heatmaps comparing NNNxxCGTG vs NNCGTG for **A.)** AhR/ARNT **or B.)** NPAS4/ARNT2. **C.)** HIF2a/ARNT or **D.)** SIM1/ARNT2. **E.)** Scatterplot comparison of Kmer affinities AhR/ARNT (y) vs NPAS4/ARNT2 (x) subset for and coloured by GTG(1-3)CGTG (red – GTGCGTG, lightblue = GTGNCGTG, green = GTGNNCGTG, darkblue = GTGNNNCGTG). **F-G.)** increased minor groove width (MGW) at NNNCGTG correlates with increased affinity (high = blue, low = Red) for **F.)** AhR/ARNT and **G.)** NPAS4/ARNT2 (left panel - all kmers, right panel – kmers subset up [T/G]CGTG). **H.)**Scatterplot comparison of Kmer affinities HIF2a/ARNT (y) vs SIM1/ARNT2 (x) subset for and coloured by GTG(1-3)CGTG (red – GTGCGTG, lightblue = GTGNCGTG, green = GTGNNCGTG, darkblue = GTGNNNCGTG). **I-J.)** increased minor groove width (MGW) at NNNCGTG correlates with increased affinity (high = blue, low = Red) for **F.)** HIF2a/ARNT and **G.)** SIM1/ARNT2 (left panel - all kmers, right panel – kmers subset up [A/G]CGTG).

Sim1 is critical for hypothalamic satiety signaling and loss of function leads to hyperphagic obesity in mice and is associated with early onset obesity and prader-willi like features in humans^23–25^. One human SIM1 early onset obesity variant, SIM1.R171H^23^, is within the PAS A -AT-Hook region and predicted to contact DNA (Fig. 3E) and shown to possess only 30% of wt SIM1 activity, but had not demonstrated whether this variant was a causal driver human monogenic obesity.

Using reporter gene assays designed to individually assess DNA binding and dimerisation (Fig. 3F and G), and interaction proteomics (Fig. 4H and Supplementary Fig. 8A-E), we found that SIM1.R171H loss of function is a result of decreased DNA binding and not altered dimerization with ARNT, ARNT2 or other interacting proteins, consistent with the PAS A loop extension mediating high affinity DNA binding. In order to investigate the biological consequence of PAS-A loop DNA binding mutant R171H and its possible contribution to hyperphagia induced obesity we generated a knock-in mouse model for SIM1.R171H (Fig. 3I-J). We found that SIM1^R171H/+^ or SIM1^R171H/R171H^ mice gained significantly (p < 10^-6^, repeated measures ANOVA) more weight on a high-fat diet over a 15 week period as compared to littermate WT controls with an associated increased food consumption (Fig. 3J and K). This showed that shape-directed distal DNA interactions made by the SIM1 PAS A loop are critical for hyperphagia induced obesity in a human obesity associated variant Sim1.R171H.

In addition to downstream shape-mediated DNA binding we also noted that nucleotides upstream of the core were associated with shape-influenced affinity relationships in bHLH-PAS TFs. By comparing the nucleotide co-dependencies in kmer affinities upstream (NNN<u>NNCGTG; f_-1_-f_-3_</u>) of the core (Fig 4. A-D) strong upstream co-dependencies in NPAS4/ARNT2 and SIM1/ARNT2 DNA binding sequences were observed that contribute to transcription factor specificity (Fig 4. E and H). In addition, AhR/ARNT and NPAS4/ARNT2 bound with similar affinity to GTG<u>CGTG</u> sequences, which appears to be shape encoded by increased MGW upstream of the core (CGTG), allowing NPAS4/ARNT2 a more flexible DNA binding motif (GTG<u>CGTG</u> or GTGT<u>CGTG</u>) as compared to AhR/ARNT (GT<u>GCGTG</u>) (Fig. 3E) and providing mechanistic detail to previous observations that NPAS4 was able to bind both TCGTG and GCGTG with high affinity^9^. NPAS4/ARNT2 appears to bind with similar to affinity to GTG<u>CGTG</u> and GTGT<u>CGTG</u> which have similar MGW – Affinity profiles indicating that NPAS4/ARNT2 may bind DNA through a shape directed mechanism (Fig. 4F-G). Similarly, SIM1/ARNT2 heterodimers bind core sequences via a shape encoded mechanism that enables specificity between TF dimers. SIM1/ARNT2 bound strongly to GTA<u>CGTG</u> and GTG<u>CGTG</u> sequences, through an associated increased MGW, whereas HIF2α/ARNT weakly bound GTG<u>CGTG</u>, contributing to additional specificity between SIM1/ARNT2 and HIF2α/ARNT DNA binding (Fig. 4H-J).

We thus find specificity that extends beyond the core is both sequence and shape encoded which allows flexibility in DNA binding. Shape encoded TF-DNA specificity for SIM1/ARNT2 and HIF2α/ARNT downstream of the core also revealed a novel AT-Hook like domain within the PAS A-loop of HIF1α and HIF2α with similarity to HMGA1 and MeCP2 AT-Hook domain amino acid sequences (Supplementary Fig. 7B). Alignment of the PAS loop domain (Supplementary Fig. 7A) of other class I bHLH-PAS transcription factors indicates that basic residues within the loop may mediate an interaction with DNA distal of the core DNA binding site as indicated by SELEX-seq and energy logo enrichment. Indeed mutations in other residues within the loop have previously been shown to lead to reduced DNA binding^16^, transactivation^23, 24, 26^ or target gene activation^27^. We also observe several other variants in HIFα and NPAS4 in the key Arg residues in the PAS A loop from clinically relevant cohorts^28^ that are likely to reduce DNA binding, providing a mechanism for observed clinical outcomes (Supplementary Fig. 7F).

Intriguingly, the presence of AT-rich selection downstream of the core sequence NNCGTG, appears to match a preference of MeCP2 binding at AT rich sequences downstream of the CmG sites^18, 22, 29^. In addition, analysis of CpH methylation sites in neurons reveals a propensity of CAC (GTG) CpH sites^30^ which also appear to have a downstream preference for A^31^ and are also preferentially recognized by MeCP2^18^. This represents an attractive mechanism by which MeCP2 may specifically target methylated bHLH-PAS motifs. Indeed, we find that bHLH-PAS transcription factor dimers are sensitive to response element methylation which appears to direct cell-type specific chromatin occupancy (Supplementary Fig 9 and 10, and **supplementary discussion**).

While we identify several key distinct mechanisms of DNA-binding selectivity within the bHLH-PAS transcription factor family, it is clear that additional mechanisms must contribute the chromatin selectivity and target gene specificity. For example, in addition to identical binding of HIF1α/ARNT and HIF2α/ARNT by SELEX seq, we also demonstrated that DNA binding affinity, hypoxic induction dynamics, stoichiometry and dimerisation strength with ARNT were unlikely to play a role in observed HIF1α vs HIF2α chromatin site selection *in vivo* (Supplementary Fig. 4A-G, see **supplementary discussion**)^5^.

In summary, we describe several important and unique features of motif recognition and DNA-protein interaction that explain interfamily specificity bHLH (MAX) vs bHLH-PAS, intrafamily specificity through core bHLH-PAS motif differences, and flanking sequences that contribute to core shared intrafamily specificity. We show that bHLH-PAS transcription factors bind DNA through a shape directed mechanism that can dictate both core flexibility and specificity as well as downstream flanking preference. We also identify novel distal PAS A loop interactions with downstream DNA sites that are important for DNA binding strength, offering explanation for the underlying cause of human hyperphagic obesity by the SIM1 R171H variant, which we recapitulated in a SIM1.R171H mouse model, and predict similar mechanisms for clinically relevant NPAS4 and HIF-2α variants.

## Methods

### Purification of bHLH-PAS heterodimers

Truncated bHLH-PAS constructs were cloned into MultiBAC baculoviral transfer plasmid pFBDM^32^ with a 6xHis-TEV leader with isothermal assembly^33^, such that each construct expressed a class I tagged transcription factor and an untagged ARNT or ARNT2 from a single baculovirus. pEFIRESpuro-hARNT-3xFlag and pEFIRESpuro-hARNT2-3xFlag, pET16b-6xHis-TEV-hAhR(1-287) (AmpR) and pAC28-hARNT(1-362) were described previously^34, 35^. Baculoviral expression and purification MultiBAC-LoxP-EYFP was generated by recombination of pUDCM-EYFP into the MultiBAC genome by Cre transposition as described^32^. With the exception of NPAS4, all truncated bHLH-PAS (A and B) heterodimers were incorporated into the MultiBAC-LoxP-EYFP baculoviral genome by pFBDM Tn7 transposition as described^32, 36^. Truncated mNPAS4 (1-329)/mARNT2 (1-481)/mnucTomato (monomeric nuclear) were recombined into MultiBAC by Tn7 transposition. Baculovirus was produced by transfection into Sf9 cells cultured in SF900III media as described^36^ and protein production monitored by eYFP or nucTomato fluorescence microscopy.

Bacterial AhR expression and purification BL21(DE3) (LysS) bacterial cells were co-transformed with pET16b-6xHis-TEV-hAhR(1-287) (AmpR)/pAC28-hARNT(1-362) (KanR). Bacteria was grown in Luria Broth to an OD600 of 0.6 and protein expressed was induced by the addition of 1mM IPTG at 16°C for 18hrs.

Insect or bacterially expressed proteins were purified by His-tag purification using hiTRAP FF columns (GE) or HisPur Resin (Thermo Scientific), followed by His tag removal by Tobacco Etch Virus (TEV) protease (in house generated) cleavage, ion-exchange chromatography, and/or size-exclusion chromatography. Protein purity was assessed by SDS-PAGE and Coomassie staining, and concentrations estimated by A280 absorbance. Purified proteins were stored in a buffer containing (20mM Hepes pH 8.0, 300mM NaCl, 5% Glycerol, 10mM DTT), and flash frozen in LiN_2_, for long term storage at −80°C. This approach left untagged heterodimeric bHLH-PAS transcription factor complexes.

Expi293 mammalian expression and purification C-terminal 3xFlag tagged full-length ARNT or ARNT2 plasmids (pEFIRES-Puro-hARNT-3xFlag or pEFIRES-Puro-hARNT2-3xFlag) were transiently transfected (200μg) into 100ml of 1.5 x 10^6^ cell/ml Expi293 cells (Life Technologies) grown in Expi293SF media using PEI Polyethylenimine (PolySciences). ∼60hrs post-transfection, Expi293 cells were harvested by centrifugation, and washed with PBS and lysed in 4ml of 20mM Hepes pH 8.0, 500mM NaCl, 1% triton-X-100, 1mM EDTA, 2xPI, 2mM NaVO_4_, 10mM Beta-Gylcerophosphate, 10mM NaF, 10% Glycerol. 75μl of Flag M2 resin used and incubated O/N at 4°C end on end, the Flag M2 resin was then washed with 5 x 1 ml lysis buffer washes, 2x 1ml 0.1% Chaps wash buffer (20mM Hepes pH 8.0, 0.1% CHAPS, 250mM NaCl, 1mM EDTA, 5% Glycerol), 2x 1ml 0.02% NP40 wash buffer (20mM Hepes pH 8.0, 0.02% NP-40, 250mM NaCl, 1mM EDTA, 5% Glycerol) and eluted with 150μl of NP40 wash buffer + 250ng/ml 3xFlag peptide end on end 1hr incubation at 4°C. Purified ARNT-3xFlag or ARNT2-3xFlag homodimers were then enriched using 100KDa mwco filters (Ambion), while removing some contaminants. The concentration and purity of ARNT-3xFlag or ARNT2-3Flag was estimated by SDS-PAGE and commassie staining with BSA standards.

### SELEX-seq

250nM of purified transcription factor and 200nM (1.5μl of 5μM of DNA library (Random 18mer or FixedCore 18/22mer)) of Fam labelled Round 0 library were incubated at room temperature in buffer containing 20mM Tris-HCl pH 8.0, 3mM MgCl2, 200-300mM NaCl, 8% Glycerol, 50 μg/ml polydI-dC, 0.2 mg/ml BSA, 5mM *β*-Me in 30μl for 20mins. DNA was extracted, amplified as described in^12^. DNA isolated from SELEX Round 0 (initial Library), Round 1 and Round 2 (FixedCore 8NCGTG10N), or Round 3 (Random 18mer) were barcoded as described in^12^ and illumina compatible adapters added by limited cycle PCR and cleaned up by PAGE purification. FixedCore 18/22mer SELEX samples from each round (Round 0, Round 1, Round 2) of were quantified, pooled and run on separate lanes of a Hiseq2500 run. Random 18mer SELEX samples (Round 0 and Round 3) were barcoded by limited cycle PCR, quantified, pooled and sequenced on a NEXTSeq500 using a single end 1×75bp High Output mode, resulting in 20-30 million reads per sample. Oligo’s used for SELEX-seq and EMSA experiments are available in Supp table 1.

Base-calling and demultiplexing was achieved with bcl2fastq, fastq files were quality filtered using Fastx toolkit^37^. In some analysis, Kmer enrichment and transcription factor binding models were generated with filtered fastq data to remove more than one “Core” binding site per read. One Core motif per read was filtered using fastq-tools (https://github.com/dcjones/fastq-tools, version 0.8) fast-grep -v function. Kmer counting and relative enrichments were analysed using the SELEX-seq R package^11, 12^ (version 1.2), transcription factor binding motif models generated using No Read Left Behind (NRLB)^13^ https://github.com/BussemakerLab/NRLB run on The University of Adelaide High Performance Computing node. Demultiplexing and trimming of primers and barcode sequences were removed using SELEX-seq or NRLB during analysis by specifying flanks and barcodes. NRLB models in GSE159989. MAX SELEX data from <u>PRJEB25690</u> EBI^13^ was also filtered for one core per line prior to re-analysis with SELEX-seq and NRLB to minimise multiple binding events per sequence. NRLB models were used to score BED using a custom R script modified from NRLBtools.

### DNAShape analysis

To investigate DNA shape contribution to the binding affinity of bHLH-PAS transcription factors we initially used shapelyzer^38^ to investigate mononucleotide NRLB model derived shape affinity relationships. We also used DNAShapeR package to analyse kmers aligned around the ‘Core’ CGTG using 14mer affinity tables from fixedCore SELEX-seq to estimate shape parameters and created affinity binned mean shape profiles to investigate shape-sequence-affinity relationships. In addition, we used NRLB dinucleotide models for HIF2a/ARNT or SIM1/ARNT2 to affinity score all 10mers downstream of a fixed upstream sequence containing the TACGTG core (HIF2a = tggAATGTG**TACGTG**NNNNNNNNNNcca, SIM1 =tggAAAGGG**TACGTG**NNNNNNNNNNcca) and shape profiles using DNAShapeR. Again, we analysed affinity binned mean shape profiles to investigate the relationship between downstream PAS A loop interactions with Affinity-DNA shape correlations. Affinity heatmap comparisons of nucleotide co-dependencies were generated using R.

### EMSA and methylC EMSA

Electrophoretic mobility shift assays were performed with different competitor DNA (ssDNA-salmon sperm DNA) conditions than SELEX-seq using Fam-labled dsDNA probes were generated by annealing upper fam labeled oligos (IDT DNA, **Supplementary table 1**). Briefly, protein DNA complexes were formed *in vitro* by incubating increasing amounts of transcription factor (0-5μg) with 10nM of EMSA probes in a buffer containing 20mM Tris pH 8.0, 250mM NaCl, 160 μg/ml ssDNA, 30 μg/ml BSA, 1.25mM MgCl_2_, 6% Glycerol, 10mM DTT. Transcription factor DNA complexes were incubated at room temperature for 30mins before separation of bound complexes by non-denaturing 5-7% PAGE. Gels were scanned using chemidoc (Biorad) with the fluorescein channel and bands intensity estimated using the imagelab software (Biorad). Relative binding of protein to different DNA probes was estimated by fraction of probe bound at a constant sub-saturating protein concentration across at least 3 independent experiments.

### Generation of SIM1.R171H humanized obesity variant knock-in mice

Mouse embryonic stem cells (C57B6/Sv12) were targeted by electroporation a SIM1 targeting construct containing exon 5 of SIM1 carrying the R171H and the loxP floxed testis specific CRE Neo selection cassette for selection in ES cells and Cre mediated removal upon germline transmission^39^. Targeting was performed essentially as described in^39^ and confirmed by PCR using MW21 F 5’ aggggcattgcaccattacag 3’, MW21 R 5’ cttgtagccaccgcaggtgaggccagc 3’ ACN F 5’ gaattcgcccttatcggcg 3’, ACN R 5’ aagctttcgcgagctcgag 3’, R MW25 5’ aaggctttggttcttaacttcc 3’. Knock-in mice were generated by blastocyst chimera generation and backcrossed onto C57/b6 background for 5 generations. Sim1 R171H genotyping was achieved with MW21/MW22/MW25 multiplexed PCR primers to detect R171H allele using KAPA Mouse Genotyping Kit.

### Mouse feeding studies

Mice were bread in accordance with The University of Adelaide laboratory animal services standard procedures and experiments approved with the animal ethics committee (approval number S-2020-027). In brief, weight gain experiments were undertaken using female mice with littermate controls from Sim1^R171H/+^ x Sim1^R171H/+^ crosses. Upon weaning at 4 weeks mice were fed ad labitum a high fat diet (HFD, SF00-219 Specialty foods) and weighed weekly for 15 weeks. Repeated measures ANOVA with Bonferroni correction for multiple comparisons was used to compare differences in weight gain between WT and R171H/+ and R171H/R171H mice on high-fat diet (0-15 weeks) from 4 weeks of age using prism. Number of mice per genotype are noted in Fig. legend. Sim1^+/+^ (n = 7) vs Sim1^R171H/+^ (n = 7) total HFD food consumed per mouse during 12 hr period (Dark) was measured using CLAMS cages (Columbus Instruments).

### Interaction proteomics

See supplementary methods.

### Mass spectrometry analysis

Peptide identification and label free quantification of mass spectrometry data was performed using MAXquant^40^ with a mass error tolerance of 20 ppm and Perseus^41^ was used for normalization, filtering and imputation. For comparison of interacting proteins between SIM1 WT and SIM1.R171H we subtracted mock purifications from immunopurified samples log transformed, cluster normalised in perseus and pearsons correlation was calculated on the label free quantification of each protein for SIM1 WT vs SIM1.R171H. To analyse relative coimmunopurification of ARNT or ARNT2 with SIM1 WT or SIM1 R171H we combined label free quantification of ARNT and ARNT2 peptides. Interacting proteins for all proteomics experiments can be found in **Supplementary Table 2**.

### ChIP-seq analysis

Bed files were used to for NRLB motif model scoring in R and a motif identification and enrichment using HOMER^42^ from NPAS4 ChIP-seq was from mouse cortical neurons depolarised with 55mM KCl (top 11,344 peaks) ^43^ (GSE21161), NPAS4, ARNT and ARNT2 Rat hippocampal neurons^7^ (GSE127793), HIF1a and HIF2a and ARNT HepG2 HK8C hypoxia treated cells^5^ (GSE120887). For HIF1a and HIF2a ChIP-seq hypoxic treated MCF7 cells^6^ (GSE28352) and AhR and ARNT ChIP-seq from TCDD treated MCF7 cells^44^ (GSE41820), were mapped to hg19 using bowtie2^45^ and peaks recalled from sequencing data using MACS2^46^. Intersections, manipulation of peaks and random sequences was achieved using bedtools^47^ or in R using Granges. Motifs from ChIP-seq peaks were identified using HOMER with the findMotifs.pl function. Receiver operator curve analysis was implemented in R with pROC^48^ comparing NRLB model derived mononucleotide scores (As described in ***SELEXseq***) of ChIP peaks or randomly selected sequences and was used to calculated AUCROC and associated errors. Linear regression of ChIP-peak scores vs motif scores was analysed and ploted as binned motif scores (0-10) vs average ChIP-peak scores in R. All statistical comparisons were analysed using R.

### Reporter assays

pGL4-Gateway-SCP1 a gift from Alexander Stark^49^ (addgene #71510) was used to clone in 6xCME by a novel rolling circle amplification and cloning procedure. Briefly, using 5’ phosphorylated CME 5’ <u>cagagcca</u>tcactgacatctgtgg**cacgta**caaatttcaatgt<u>ggaaggctg</u> 3’ and rolling circle primers RCA1 (KpnI) 5’ atatggtacctctgcagccttc 3’ and RCA2 (BglII) 5’ atatagatctgctgcagagcca 3’. Briefly, 10pmol of template oligo was cyclized and ligated using CircLigase II ssDNA Ligase (Lucigen) supplemented with 2.5mM MnCl_2_, 1M Betaine, 1x CircLigase II Buffer and 100U of CircLigase II enzyme in a 20μl reaction incubated at 60°C for 1 hr and 80°C for 15mins. The reactions were then purified by phenol:chloroform clean up and used in Rolling Circle Amplification (RCA) reactions. RCA reactions were performed with 20ng of circular ssDNA, 5μl of 10x BstPol II buffer (NEB), 1.5μl of 10mM dNTPs, 1μl of T4 bacteriophage protein (NEB, 5ng/μl), 3μl of DMSO, 1μl of RCA1 (60μM) and RCA2 (60μM) and 0.8μl of BstII Polymerase (NEB, 8U/μl) in a 50μl reaction. The RCA was then performed at 65°C for 90mins followed by 55°C for 120mins, the RCA reactions were then separated by agarose gel electrophoresis and repeat lengths of interest cloned into pGL4-Scp1 and sequence verified.

Expression constructs containing N-Terminal class I bHLH-PAS transcription factors (HIF1a (1-375), HIF2a(1-353), SIM1 (1-438), SIM1.R171H (1-438) fused to the VP64-p65-RTA (VPR) activation domain^50^ were cloned into pEFIRESpuro expression plasmid by isothermal assembly. N-terminal truncated ARNT (1-503)-2Myc, ARNT2(1-455) −2Myc, Gal4DBD-ARNT (1-503)-2Myc were also cloned into pEFIRESpuro expression plasmids. Reporter assays were performed in 96 well white Griener μclear plates (655094) by seeding 1 x 10^4^ HEK293T cells/well. The following day, cells were transfected with 100ng of pG5e1b^51^ or pGL4-scp1-6xCME reporter plasmid, 0.5ng pCI-RL (Renilla) plasmid and 25ng of each expression plasmid or a empty vector with PEI. 48hrs after transfection firely luciferase was assayed in plate using a LARII (Promega) and measured on a GloMax luminometer.

### CRISPR knock-in of tags to endogenous HIF1a and HIF2a

CRISPR targeting constructs clones targeting adjacent to the endogenous HIF1a and HIF2a stop codons were cloned into px330 by ligating annealed and phosphorylated oligos with BbsI digested px330, using hHIF1a sgRNA upper 5’ caccgTGAAGAATTACTCAGAGCTT 3’, hHIF1a sgRNA lower 5’ aaacAAGCTCTGAGTAATTCTTCAc 3’ or hHIF2a CTD sgRNA upper 5’ caccgCCTCCTCAGAGCCCTGGACC 3’, hHIF2a CTD sgRNA lower 5’ aaacGGTCCAGGGCTCTGAGGAGGc 3’. Knock-in of HA-3xFlag epitopes into the endogenous HIF1a or HIF2a locus in HepG2 cells was achieved by transfection with 0.625 μg of pNSEN, 0.625 μg of pEFIRES-puro6, 2.5μg of px330-sgHIFa CTD, and 1.25μg of ssDNA HDR template oligo containing flanking homology to CRISPR targeting site the tag insertion and a PAM mutant into ∼0.8×10^6^ cells using polyethylenimine (PEI, 3:1). Forty-eight hours after transfection, the medium was removed from cells and replaced with fresh medium supplemented with 2 μg/ml puromycin for 48 hrs and the cell medium was changed to fresh medium without puromycin. Forty-eight hours later cells were seeded by limiting dilution into 96-well plates such that an average of 0.5 cells/well were present. Correct integration of the tags into the endogenous loci was identified by PCR screening using HIF1a gDNA screen F 5’ ggcaatcaatggatgaaagtggatt 3’, HIF1a gDNA screen R 5’ gctactgcaatgcaatggtttaaat 3’, and HIF2a gDNA screen F 5’ taccaacccttctttcaggcatggc 3’, HIF2a gDNA screen R 5’ gcttggtgacctgggcaagtctgc 3’ and positive colonies reisolated as single colonies by limiting dilution. Isolated HIF1a and HIF2a tag insertions were confirmed by PCR, sanger sequencing and western blotting.

HepG2 cells were grown in normoxia or <1% oxygen and 5% CO_2_ using a hypoxia workstation for 4 or 16hrs. Cells were then washed with PBS prior to lysis with whole cell extract buffer (20mM Hepes pH 8.0, 420mM NaCl_2_, 0.5% Igepal, 0.2mM EDTA, 1.5mM MgCl_2_, 25% Glycerol) supplemented with 2mM DTT and 1x protease inhibitors. Extracts were run on 7.5% SDS-PAGE gels and transferred to nitrocellulose prior to western blot detection with anti-HA (HIFa, HA11 – 16B12; MMS-101R), anti-*β*-tubulin (Biorad) and species specific HRP conjugated secondary antibodies (Peirce).

### MethylC-seq Analysis

See supplementary methods

### Structural Analysis of bHLH-PAS heterodimer DNA binding

#### bHLH-PAS factor homology/PAS A loop modelling in ICM-Pro (*Molsoft ICM 3.8-6a*)

Homology models based on the mouse HIF2a:ARNT HRE DNA crystal structure (PDB: 4ZPK) were constructed in ICM-Pro ^16, 52^. Peptide sequences for human ARNT (UniProt ID: P27540), ARNT2 (UniProt ID: Q9HBZ2), SIM1 (UniProt ID: P81133) and NPAS4 (UniProt ID: Q8IUM7) were prepared in FASTA format and read directly into the ICM workspace. Sequence-template alignments were made for each model to the corresponding reference chain e.g. SIM1 peptide sequence to the HIF2a protein structure. Homology models were constructed using Model Builder and subsequently refined ^53^. Double stranded DNA element models (SIM1: TGGAAAGGGTACGTGACCCGCTGCACCA; NPAS4_di: TGGAAATGGGTCGTGACCCAGGATTCCA) were built in PyMOL and refined prior to alignment by carbon-α atoms to HRE DNA. Protein/dsDNA homology models were merged and refined.

Modelling of the PAS A loop to investigate potential DNA contacts was performed by *ab initio* interactive loop modelling of the local environment, within the internal coordinate mechanisms force field (ICMFF) ^54^. Modelling of the HIF2a:ARNT (4ZPK) PAS A loop on HRE and SIM1 dsDNA was performed as control, as well as modelling the HRE for all homology models. A list of protein-DNA models and oligo sequences are available in (Supplementary table 3). Structural comparisons were made in pymol or chimera and all figures made using chimera.

Analysis of protein DNA contacts was performed using DNAproDB^55^ and manually annotated onto DNA sequences or models.

#### Statistical information

Statistical comparisons were implemented using R or Prism and indicated in the main text or figure legends.

## Data availability

SELEX sequencing data for fixed core and random 18mer data sets are available GSE159989, Kmer Affinity tables are also available at GSE159989.

## Code availability

Code used here was implemented predominantly in R. Code and NRLB models used for scoring bed files will be made available at https://github.com/BeeSting-pgm/TF_BED_Score.

## Acknowledgements

The Authors wish to acknowledge the Aboriginal Kaurna people, the original custodians of the Adelaide Plains and the land on which the experiments were performed at the University of Adelaide Campus. We thank Joel Mackay (The University of Sydney) for help in establishing the Expi293 culture system. This work was supported with supercomputing resources provided by the Phoenix HPC service at the University of Adelaide.

## Author Contributions

D.C.B and M.L.W. conceived the project and wrote the manuscript. D.C.B performed all experiments and analyzed data. All code used to analyse SELEX-seq, DNAShape and ChIP-seq was written and executed by D.C.B. J.B. analysed methylCpG sequencing data. D.C.B and M.L.W designed and generated ES targeting strategy, ES line generation, and mouse model generation. D.C.B and A.S., R.F., G.M. performed mouse experiments. D.C.B., D.M., and J.B. analysed structures and structural models.

## Materials & Correspondence

Correspondence and requests for materials should be addressed to D.C.B.

## Competing interests

No competing interests to be declared by the authors as a part of this work.

## Supplementary Information

### General background for bHLH/PAS Transcription Factors

bHLH-PAS TF family members perform distinct, non-overlapping biological roles and regulate specific target genes ^1, 2^. For example:

1) The hypoxic regulated transcription factors HIF1a and HIF2a are coexpressed in many cell types and tissues but display distinct transcription factor (HIF1a – promoter bound HIF2a – Enhancer bound) occupancy patterns and regulate overlapping and distinct genes^3–5^.

2) Several of the bHLH-PAS transcription factors display cell type specific target gene expression^4^, eg NPAS4 regulates different target genes in inhibitory *(Frmpd3, Cort, Npr3, and Rerg)* and excitatory neurons *(BDNF, Csrnp1 and Nrp1)* to execute opposing synapse functions^6^.

3) Furthermore, SIM1 and HIF1a/HIF2a have been proposed to compete for the same response element but perform distinct biological functions and do not appear to regulate each other’s target gene programs. For example, SIM1 has been shown to be definitive marker of paraventricular hypothalamic neurons and a master regulator of appetite control^7–9^. Mutations in Sim1 have also been shown to be associated with early onset hyperphagic obesity in humans although the mechanism of loss of function is unknown for the majority of these mutations.

bHLH-PAS TF’s bind NNCGTG (CACG – E-box-like) DNA sequences as heterodimers, containing one Class I (HIF1a, HIF2a, AhR, SIM1, NPAS1, NPAS3, NPAS4) protein with one Class II (ARNT or ARNT2) partner. The PAS domains restrict heterodimeric partner selection to within the bHLH-PAS subfamily and strengthen DNA binding and heterodimerisation. In addition, the PAS domains have been proposed to contact DNA outside the Core NNCGTG motif conveying additional DNA specificity^10^ or strength^11–13^. PAS domains have also been shown to mediate protein-protein interaction with co-activators or other transcription factors and PAS domain swap experiments in drosophila homologs is sufficient to switch target gene expression^10^.

Taken together the bHLH-PAS domain transcription factors are a unique bHLH sub-family for which mechanisms that define target gene selection and cell specific functions remain opaque.

Through comparative profiling of inherent DNA binding specificities *in vitro*, an interplay between DNA shape and sequence selectivity, and *in vivo* determinants of DNA binding we reveal that DNA binding specificity and chromatin selectivity is encoded by multiple mechanisms.

## Supplementary Results

### Information for Supplementary Figure 1

To answer questions surrounding bHLH-PAS transcription factor selectivity (between heterodimers for chromatin) and specificity (inherent DNA binding affinities) we profiled hierarchies of DNA sequences bound by the most distinctive members of the family (Supplementary Fig.1A).

bHLH-PAS transcription factors prototypically form heterodimer pairs with either ARNT or ARNT2. To facilitate profiling of the DNA response elements of the bHLH-PAS transcription factor family we purified full length (ARNT or ARNT2) or N-terminal truncated (bHLH-PAS) dimers from mammalian Expi293 cells (ARNT or ARNT2), insect baculovirus infected Sf9 cells (HIF1a/ARNT, HIF2a/ARNT, SIM1/ARNT2, NPAS4/ARNT2) or bacteria (AhR/ARNT) (Supplementary Fig. 1B). Dimer pairs were selected based on predicted diversity of DNA binding response elements and the most probable endogenous complexes^2^. For example, ubiquitously expressed HIFa and AhR have been shown to act in heterodimeric complexes with ARNT, whereas NPAS4 and SIM1 have been shown to partner ARNT2 in neuronal cells. ARNT and ARNT2 have also been shown to form homodimers, which we also analysed, despite the *in vivo* significance of these homodimers remaining unknown.

*Side note: CLOCK (and related NPAS2) and BMAL are bHLH/PAS family members that appear to predominantly heterodimerise with each other and not with other bHLH-PAS transcription factors (ARNT or other class I transcription factors). In addition, CLOCK/BMAL heterodimers bind to canonical E-Box elements and not E-Box-like elements, and adopt a fundamentally different heterodimeric interface of contacts compared to other bHLH-PAS family members^11, 14^, thus were not studied here as they were considered as mechanistically separate to ARNT containing heterodimers.*

**Supplementary Figure 1.**
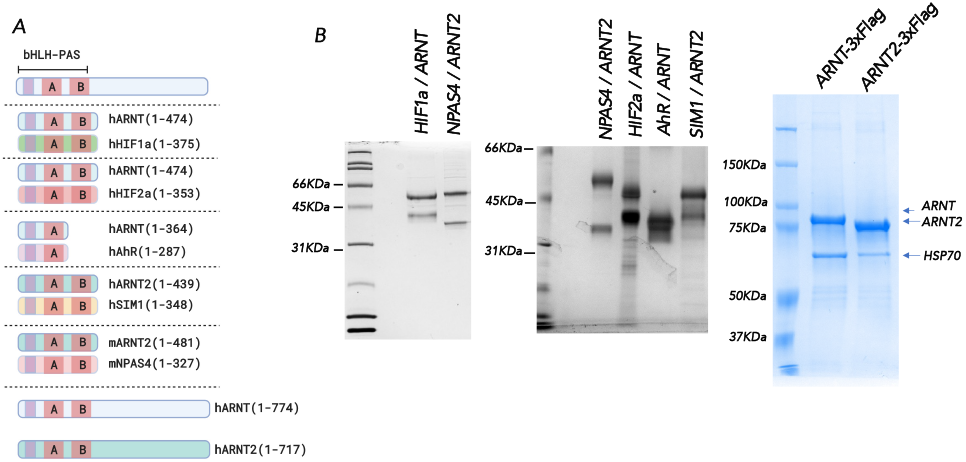
Transcription Factor truncations and proteins used in SELEX-seq. **A.)** Schematic diagram of the different bHLH-PAS transcription factor constructs used in SELEX-seq and *in vitro* DNA binding analyses. **B.)** SDS-PAGE gel analysis of purified dimeric bHLH-PAS proteins used in this study.

### Information for Supplementary Figure 2

#### Sub-Classification of bHLH-PAS transcription factors by response element binding

We used the proteins displayed in Supplementary Figure 1 in SELEX-seq experiments by incubating the heterodimers with random (18N) or FixedCore (8NCGTG10N) dsDNA libraries (Figure 1A). Following 1-3 rounds of high-throughput sequencing we analysed enriched DNA binding sites by SELEX-seq^15, 16^ R package and NRLB energy modeling^17^.

We generated affinity tables for 10mers (random 18mer library) or 12mers (18/22mer fixed core) and compared the selection of the nucleotide directly upstream of the Core CGTG (i.e NCGTG). We found strong concordance between round 1 or round 2 affinities generated for fixed core 12mers affinities (Supplementary Fig 2A; HIF2a/ARNT r1 v r2 *R^2^= 0.86*, SIM1/ARNT2 r1 v r2 *R^2^= 0.89*, AhR/ARNT r1 v r2 *R^2^= 0.86*, NPAS4/ARNT2 r1 v r2 *R^2^= 0.89*). In addition, we also observed that both library selection strategies strongly enriched for consistent transcription factor specific NCGTG response elements (Supplementary Fig. 2B-D). Consistent with previous reports we found that ARNT, ARNT2, HIF1a, HIF2a and SIM1 selectively bind ACGTG containing elements whereas NPAS4 (TCGTG) and AhR (GCGTG) bind distinct core elements (Supplementary Fig. 2E-F). As expected ACGTG binding bHLH-PAS transcription factors can be further distinguished by the nucleotide upstream of the ACGTG (NACGTG). ARNT and ARNT2 highest affinity sites select for CACGTG containing response elements, whereas HIF1a/ARNT, HIF2a/ARNT and SIM1/ARNT2 complexes select for TACGTG containing response elements, NPAS4/ARNT2 select for GTCGTG containing response elements and AhR/ARNT selecting for TTGCGTG containing response elements. These NNCGTG affinities are also sufficient to explain site preference between almost all bHLH PAS heterodimers (Supplementary Fig. 3 A-F), the major exception being a lack of distinction between HIFa/ARNT and SIM1/ARNT2.

**Supplementary Figure 2.**
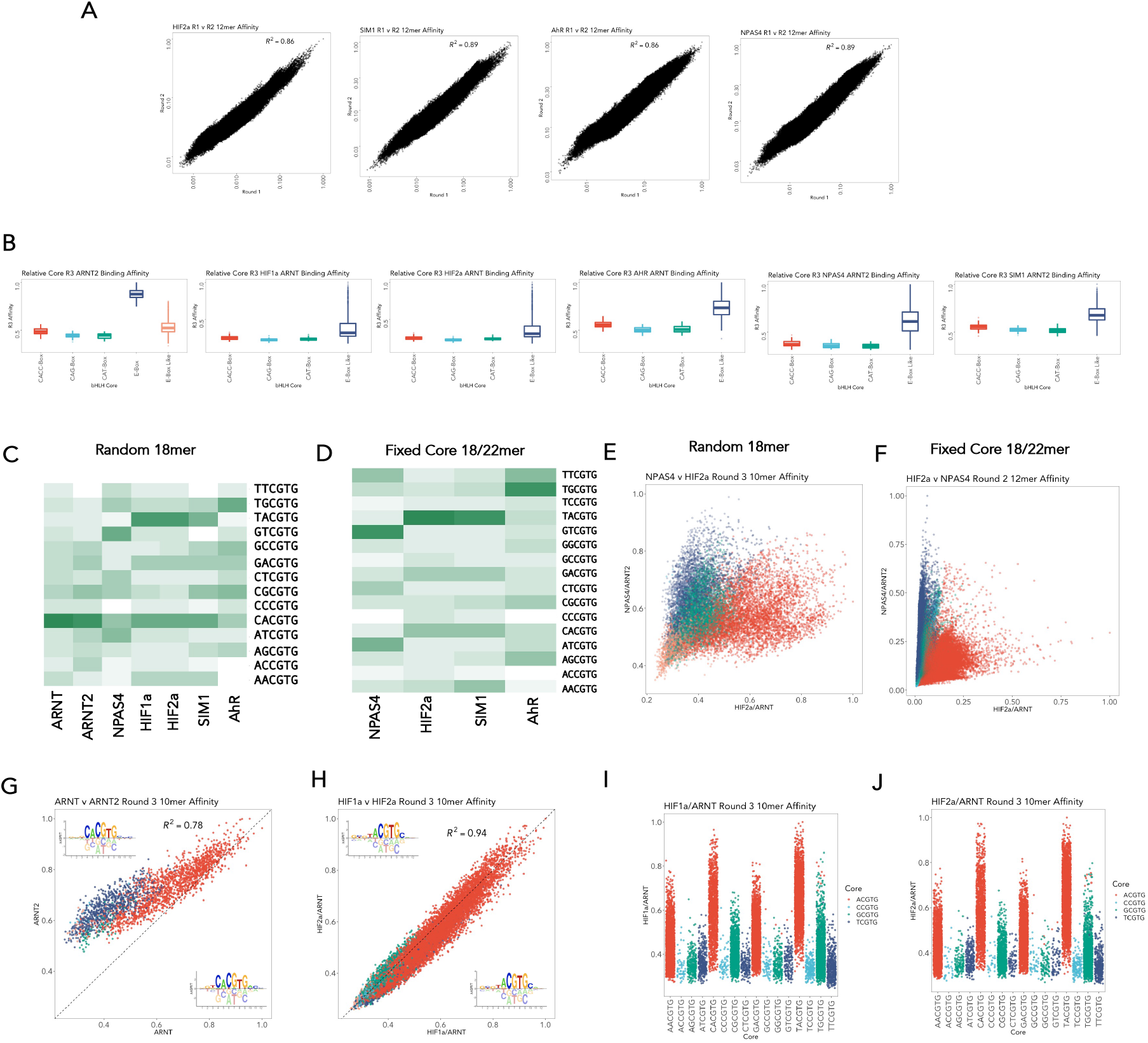
Comparison SELEX-seq analysis strategies and Core DNA binding specificity of the bHLH-PAS transcription factors. **A.)** Comparison of the relative affinity of 12mer affinities from Round 1 or Round 2 of FixedCore 18/22mer SELEX-seq. Log_10_ 12mer affinities generated from SELEX-seq analysis were plotted comparing Round 1 (x-axis) and Round 2 (y-axis) for HIF2α/ARNT, SIM1/ARNT2, AhR/ARNT or NPAS4/ARNT2. correlation between R1 and R2 was assessed by coefficient of determination r^2^. **B.)** Round 3 Kmer Affinity boxplots for each transcription factor on different bHLH motifs. **C.)** and **D.)** Heatmaps comparing relative 10mer affinities for the labeled bHLH-PAS transcription factor complexes on each core NNCGTG DNA binding site generated using **E.)** Random 18mer library SELEX strategy (Round 3) or **F.)** the FixedCore 18/22mer SELEX strategy (Round 2). **E.)** and **F.)** Differential bHLH-PAS DNA binding specificity is encoded by distinct core NCGTG sequences (Red = ACGTG, Blue = TCGTG, green = GCGTG, pink = CCGTG). Comparison of Relative Affinities of NPAS4/ARNT2 vs HIF2α/ARNT to **E.)** 10mers (Random 18mer library SELEX strategy (Round 3)) or **F.)** 12mers (FixedCore 18/22mer SELEX strategy (Round 2)). Comparison of Relative 10mers Kmer Affinities (Random 18mer library SELEX strategy (Round 3)) (Inset 12mer NRLB models) **G.)** ARNT vs ARNT2, r^2^ = 0.78 **H.)** HIF1α/ARNT vs HIF2α/ARNT2, r^2^ = 0.94. Relative Core NNCGTG 10mer Affinities of each probe for **I)** HIF1α/ARNT or **J.)** HIF2α/ARNT (Random 18mer library SELEX strategy (Round 3)).

### Information for Supplementary Figures 2 and 4

#### Analysing Isoform specificity: ARNT Vs ARNT2 and HIF1a Vs HIF2a

ARNT and ARNT2 have been shown to have some distinct biological functions despite significant overlapping expression patterns in neurons and other tissues^2^. We generated NRLB models for ARNT and ARNT2 (Supplementary Fig 2G) and compared 10mer affinities for ARNT and ARNT2 DNA binding and found a high degree of correlation between affinities (Supplementary Fig 2G **- *R^2^=0.78***). Although ARNT appears to be highly restricted to CACGTG containing response elements ARNT2 may be able to bind to a more flexible consensus (NACGTG or CGCGTG albeit with much lower affinity than the core CACGTG).

HIF1α and HIF2α display overlapping and distinct expressing patterns as well as biological functions in the response to low oxygen adaptation^18^. These disparate functions are proposed to be mediated by the distinct HIF1α or HIF2α dependent target gene expression, for example HIF1a regulates metabolic genes whereas HIF2α regulates genes involved in erythropoiesis and iron homeostasis^18^. HIF1α and HIF2α also show distinct genomic occupancy within the same cell types, with HIF1α binding more prevalently to promoter regions whereas HIF2α binding more prevalently to enhancers ^3, 4, 19, 20^. Low resolution analysis of DNA binding specificity from ChIP (RCGTG (R = A or G)) has not sufficiently addressed whether inherently different HIF1α and HIF2α DNA binding specificities might explain genomic site selection preferences. To address this we compared the DNA binding specificity of HIF1α/ARNT and HIF2α/ARNT 10mer affinity tables (Supplementary Fig 2I-J) finding a near identical DNA binding specificity (*R^2^ = 0.94*). As expected HIF1α and HIF2α bound strongly to ACGTG containing cores and less efficiently to GCGTG containing cores, most efficiently binding TACGTG (Supplementary Fig 2D, 2I-J). We also found that HIF1α/ARNT and HIF2α/ARNT bound with similar affinity (HIF1α ∼2 fold higher affinity to HIF2α) to TACGTG containing probes in EMSA analyses, consistent with previous reports^11^, and indicating that the HIF1α/ARNT and HIF2α/ARNT *in vitro* specificity and affinity is near identical and is unlikely to contribute to HIFa genome binding selection (Supplementary Fig 4A-C). We hypothesized that given that there are both overlapping and distinct HIF1α and HIF2α chromatin binding sites that sequential loading or competition for HIF binding sites may be influenced by the hypoxic induction dynamics or stoichiometry of HIF1α or HIF2α. Therefore, we generated C-terminally HA-3xFlag (HF) tagged heterozygous HIF1α or HIF2α knock-in HepG2 cells using CRISPR/Cas9 mediated homologous directed repair (Supplementary Fig. 4D). We did not observe isoform specific differences in the dynamics of hypoxic induction at <1% O_2_, with both HIF1α and HIF2α displaying hypoxic protein induction at 4 and 16hrs in HepG2 (2 x HIF1α.HF lines and 1 x HIF2α.HF (Supplementary Fig. 4E). However we observed large differences in the relative stoichiometry of HIF1a and HIF2a in HepG2 cells (Supplementary Fig. 4E). This suggests that sequential loading of chromatin by HIF1a and HIF2a is not a mechanism for chromatin site selection and that HIF1a is able unable to occupy HIF2a enhancer sites, even when in significant excess to HIF2a. This is also supported by recent evidence showing that HIF1α and HIF2α are unable to occupy each other’s preferential binding site even if the other isoform is removed^4^. Next, we developed a two-hybrid approach to assess *in vivo* dimerization strength between bHLH-PAS heterodimers. In this system we retained endogenous dimerisation domains of ARNT but removed transactivation domains and N-terminally tagged the protein with the Gal4 DNA binding domain. HIF1α or HIF2α bHLH/PAS domains were then cloned in frame with the strong transactivation domain VPR^21^ and expressed with Gal4-ARNT fusion proteins and a Gal4 responsive reporter gene in HEK293T cells to assess dimer strength (Supplementary Fig. 5F). We found that Gal4DBD-ARNT (1-503) (*Δ*TAD) was unable to activate the reporter but co-expression of HIF1α(1-375)-VPR or HIF2α(1-353)-VPR strongly activated the reporter (Supplementary Fig. 4F). Both HIF1α and HIF2α with Gal4-ARNT activated the reporter to a similar level indicating that dimerisation strength was similar between HIF1α and HIF2α. While *in vitro* DNA binding specificity and affinity for the HRE was similar between HIF1α and HIF2α we reasoned that *in vivo* DNA binding affinity or specificity may differ. Therefore, we used ARNT (1-503) (*Δ*TAD) and ARNT2 (1-455) (*Δ*TAD) co expressed with HIF1α(1-375)-VPR or HIF2α(1-353)-VPR on the bHLH-PAS reporter 6xCME (TACGTG). Surprisingly, we found that HIF1a-VPR more strongly activated that reporter than HIF2a-VPR despite similar dimerisation (Supplementary Fig. 4G). This could be either explained by differences DNA binding affinity between *in vitro* and *in vivo* experiments or differential protein-protein interactions mediated through the N-terminal domains.

**Supplementary Figure 3.**
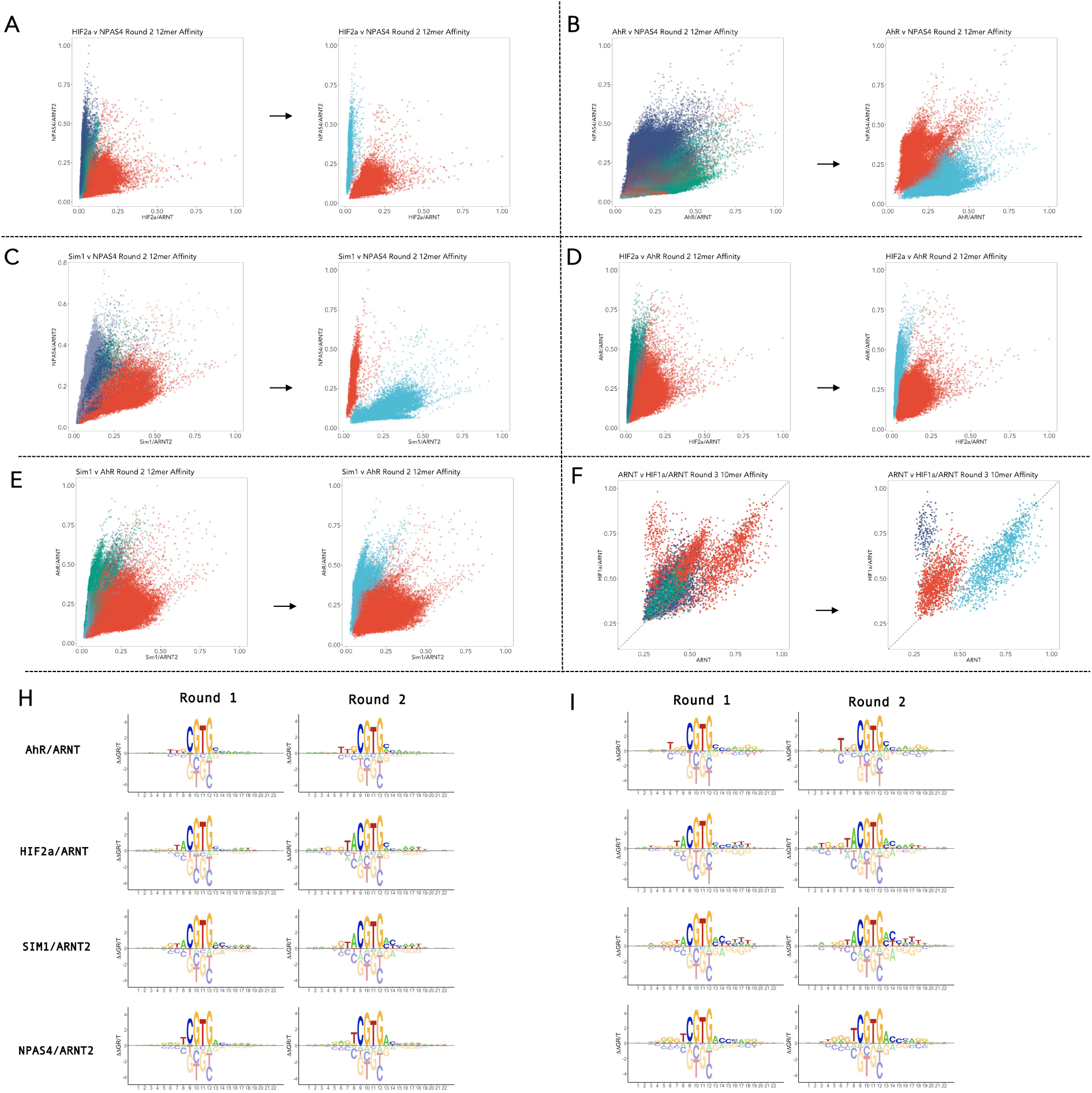
Core encoded specificity and energy models of the bHLH-PAS transcription factor family. **A-F.)** Kmer Affinity scatter plots Core specificity colour by NCGTG (left panel) or the most divergent NNCGTG right panel. **A.)** NPAS4/ARNT2 v HIF2α/ARNT (Round 2 12mers - fixedCore 18/22mer) left panel (ACGTG - red, TCGTG – darkblue), right panel (GTCGTG - blue, TACGTG - red). **B.)** NPAS4/ARNT2 v AhR/ARNT (Round 2 12mers - fixedCore 18/22mer) left panel (GCGTG - darkblue, TCGTG – green), right panel (GTCGTG - red, TGCGTG - blue). **C.)** SIM1/ARNT2 v NPAS4/ARNT2 (Round 2 12mers - fixedCore 18/22mer) left panel (TCGTG - red, ACGTG – mauve), right panel (GTCGTG - red, TACGTG - blue). **D.)** HIF2α/ARNT v AhR/ARNT (Round 2 12mers - fixedCore 18/22mer) left panel (TCGTG - green, ACGTG – red), right panel (TACGTG - blue, TGCGTG - red). **E.)** SIM1/ARNT2 v AhR/ARNT (Round 2 12mers - fixedCore 18/22mer) left panel (GCGTG - green, ACGTG – red), right panel (TGCGTG - blue, TACGTG - red). **F.)** ARNT/ARNT v HIF1α/ARNT (Round 3 10mers - random 18mers) left panel (GCGTG - darkblue, TCGTG – green, ACGTG – red), right panel (CACGTG - lightblue, TACGTG – darkblue, GACGTG - red). Energy Logos generated from Round 1 vs Round 2 FixedCore **H.)** mononucleotide or **I.)** dinucleotide models. NRLB non-symmetrical models were generated by modeling on SELEX-seq filtered data (one central CGTG per read) from Round 1 (left panel) or Round 2 (right panel) for AhR/ARNT, HIF2α/ARNT, SIM1/ARNT2 and NPAS4/ARNT2.

**Supplementary Figure 4.**
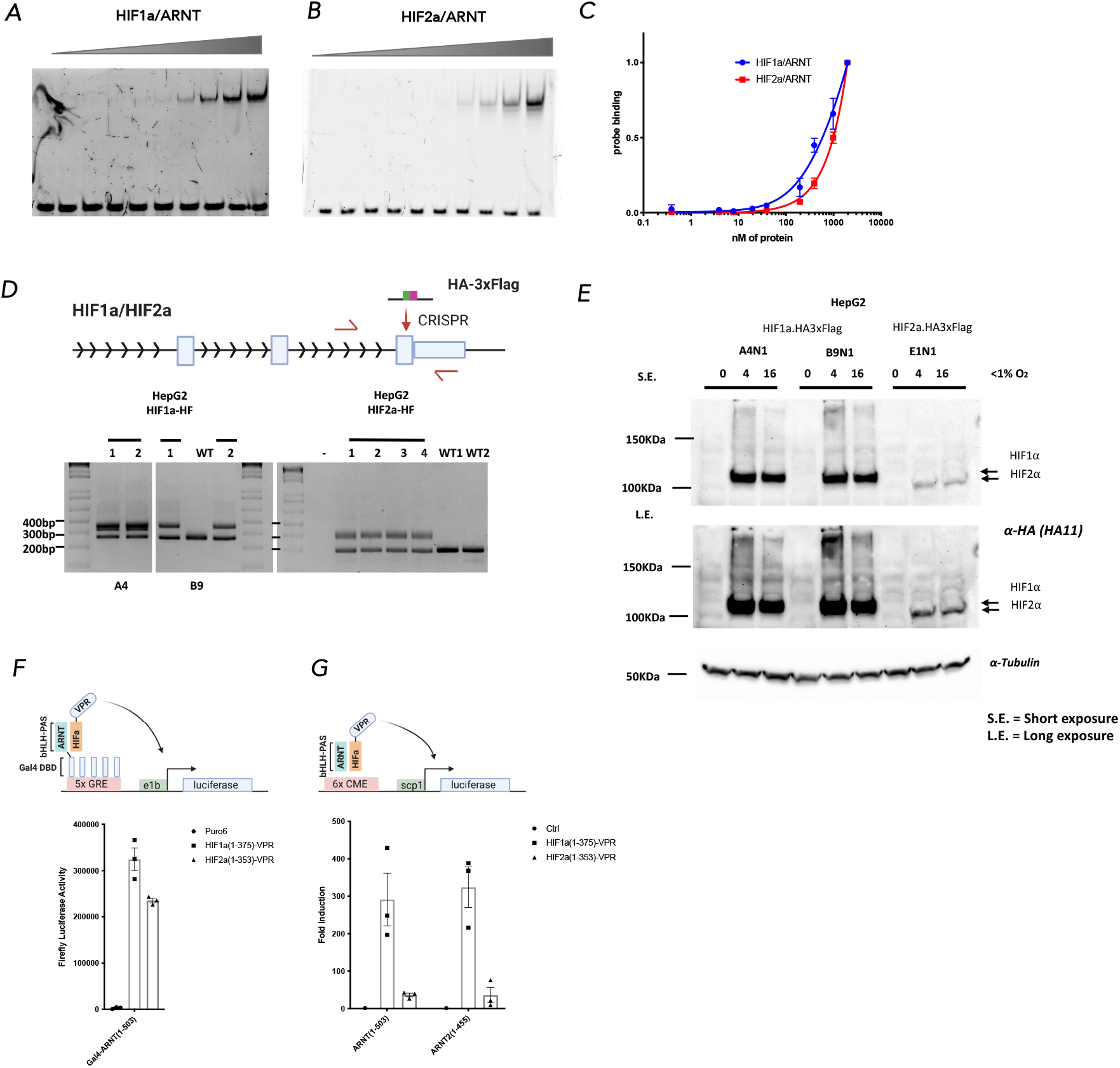
Characteristics of HIF1α vs HIF2α DNA-binding, dimerization and stoichiometry. A.) and B.) DNA binding Affinity of HIF1α/ARNT and HIF2α/ARNT was assessed by EMSA. Increasing amounts of A.) HIF1α/ARNT or B.) HIF2α/ARNT heterodimer were incubated with a FAM labeled HRE probe (AGGCTGCGTACGTGCGGGTCGT). C.) Relative HRE probe binding from gel shift experiments (n = 3) shows similar DNA binding affinity for HIF2α/ARNT (red) or HIF1α/ARNT (blue). D.) Schematic diagram (upper panel) of CRISPR homology directed repair (HDR) strategy to knock-in HA-3xFlag tag into endogenous HIF1a or HIF2α locus. CRISPR guide sgRNA was designed to cut near the endogenous stop codon of HIF1α or HIF2α in HepG2 cells, and an oligo template containing flanking homology to HIF1α or HIF2α and a single HA tag and 3xFlag tags was provided as a HDR template. Cells monoclone’s were isolated by two rounds of limiting dilution and genomic PCR screening for HA-Flag insertion (Using primers flanking the insertion site (red)), followed by sanger sequencing to confirm tag insertion. E.) Western blot of HIF1α and HIF2α tagged HepG2 monoclonal cell lines. HepG2 cells were incubated in 0.5-1% O2 chamber for 4 or 16 hrs prior to protein extraction SDS-PAGE gel electrophoresis and western blotting with anti-HA or anti-Tubulin antibodies. F.) and G.) To assess in vivo dimerization and DNA binding differences between HIF1α and HIF2α we removed HIF1α or HIF2α transactivation domains (Δ TAD) and fused them to the strong heterologous transactivation domain VPR. HIFα-VPR fusion proteins were expressed in HEK293T cells together with F.) Gal4-DBD-ARNT (bHLH-PAS (1-503), Δ TAD) and the Gal4 responsive luciferase construct or G.) ARNT (bHLH-PAS (1-503), Δ TAD) or ARNT2 (bHLH-PAS (1-455), Δ TAD) and the bHLH-PAS responsive luciferase construct (6xCME, tgaaatttgTACGTGccacagatg). Firefly Luciferase activity was measured 48hrs after transfection and data is mean ± SEM of 3 experiments.

### Information for Supplementary Figures 3 and 5

#### Correlation between SELEX-seq derived DNA-binding and ChIP-seq derived DNA binding Affinity

Next, we generated energy logos from the NRLB models for all SELEX-seq data (Supplementary Fig. 3H-I **and** Supplementary Fig. 5B). Both mononucleotide and dinucleotide models derived from either round 1 or 2 of the FixedCore library approach were similar and revealed extensive flanking nucleotide preferences. Consistent with previous validation of the accuracy of NRLB modeling approach^17^ we confirmed the ability of models to accurately predict DNA binding.

Firstly, we validated flanking nucleotide contribution to binding affinity by EMSA of NPAS4/ARNT2 heterodimers on the top mononucleotide 22mer sequence derived from NRLB models compared to flanking variants of this sequence (Supplementary Fig. 5A). We found that variants upstream or downstream of the invariant core (GTCGTG) contributed to binding affinity (Supplementary Fig. 5A). We found a linear relationship between predicted binding affinity and comparative EMSA probe binding (Supplementary Fig. 5A**; r^2^=0.94**), indicating that models were able to predict flanking nucleotide contribution to binding affinity. Next, we qualitatively compared motifs derived from ChIP-seq experiments (HOMER motifs) to NRLB logos from both library strategies. We found that while ChIP derived motifs generally had lower information content than SELEX motifs but the upstream core (NCGTG or NNCGTG) nucleotide preferences matched well with NRLB logos (Supplementary Fig. 5B). Next, we compared the ability bHLH-PAS DNA binding models to predict ChIP-seq peaks^3, 4, 22–24^ by energy model-based scoring of ChIP peaks as compared to random genomic regions using area under receiver operator curves (AUROC) to assess model performance. Comparison of the maximal binding score vs AUC of all scored binding events at each peak found little difference in the ability to identify NPAS4 ChIP-peaks^23^ (AUROCmax = 0.69 vs AUCROCauc = 0.67) and we used maximal score per peak for subsequent analysis.

We found that NPAS4/ARNT2 models accurately predict NPAS4 occupancy at NPAS4, NPAS4/ARNT or NPAS4/ARNT2 peaks, and performed better than ARNT or ARNT2 models (Supplementary Fig.5C; rat PTX AUROC = 0.69-0.72). We also found that models were better at discriminating strongly bound peaks vs weakly bound peaks (Supplementary Fig.5C, rat PTX; AUROC = 0.68 vs 0.92, mouse KCl; AUROC 0.63 v 0.79). We reasoned that DNA binding affinity would result in increased ChIP-peak intensity. As such, we also compared the models affinity scores to *in vivo* ‘affinity’ using linear regression by comparing mean ChIP-peak intensity vs binned (1-10) predicted motif affinity. We found a highly significant relationship between NPAS4 affinity scores and NPAS4 ChIP -peak intensity (mouse and rat p < 2 x 10^-16^, Supplementary Fig. 5D-E-F). We also find larger linear regression coefficient (Supplementary Fig. 5D; **inset**) for NPAS4 TF-ChIP peak (NPAS4, NPAS4/ARNT, NPAS4/ARNT2) linear regression as compared to ARNT or ARNT model-ChIP peak (NPAS4, NPAS4/ARNT, NPAS4/ARNT2) linear indicting NPAS4 selectivity for NPAS4 ChIP peaks.

We also compared the ability of ARNT, HIF1α/ARNT or HIF2α/ARNT (random 18mer and 18/22 FixedCore) derived models to accurately discriminate ChIP bound peaks in HepG2. While all models were able to accurately identify ChIP-peaks HIFα/ARNT models performed best at predicting HIFα/ARNT shared peaks (AUROC > 0.9 at HIFα/ARNT peaks) (Supplementary Fig. 5F). HIF1α, HIF2α and AhR MCF7 ChIP-seq comparison also confirmed model prediction of ChIP peak intensity (Supplementary Fig. 5H-J; HIF1α ChIP, HIFα model AUROC = 0.75, HIF2a ChIP, HIFα model AUROC = 0.68, AhR/ARNT ChIP-peaks, AhR model AUROC = 0.70). Taken together this validates the ability of *in vitro* generated TF models to accurately predict and quantify both *in vitro* and *in vivo* binding.

**Supplementary Figure 5.**
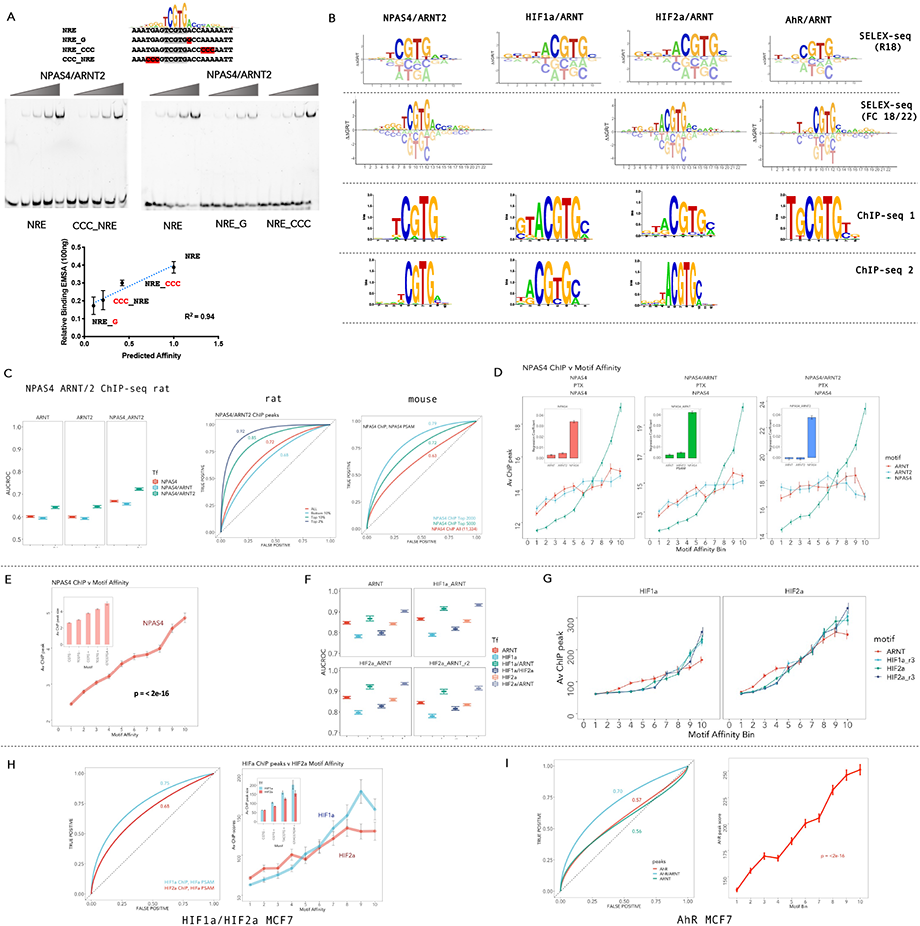
Validation of DNA binging Energy Models. **A.)** NRLB Energy Logo and EMSA probe sequences used to compare flanking nucleotide contribution to DNA binding. EMSA of increasing amounts of NPAS4/ARNT2 bound to variant Fam-labled dsDNA probes. **(Bottom panel)** quantification of EMSA binding at submaximal (100ng) EMSA binding vs predicted Affinity from NRLB mononucleotide model. Mean (±SEM) of 3 independent experiments. **B.)** representative models generated by SELEX-seq were compared to models predicted by HOMER from ChIP-seq data. **C.) (left panel)** Area under receiver operator curves (AUROC) (±SD) for the ability of NRLB DNA binding model for ARNT, ARNT2 or NPAS4 to predict NPAS4, NPAS4/ARNT, or NPAS4/ARNT2 ChIP-peaks vs size matched random sequences. NRLB models were used to score ChIP-peaks or randomly selected size matched regions to compare the ability to identify true positive transcription factor binding sites. **(right panel)** Receiver operator curves comparing the model-based prediction of NPAS4 occupied sites for rat or mouse ChIP-seq subset by peak score (i.e top 10% of sites = top 10% of peak scores for rat, i.e top 2000 or 5000 of sites by peak score for mouse). The AUROC is indicated on each corresponding line. **D.)** NPAS4, NPAS4/ARNT or NPAS4/ARNT2 ChIP-peak DNA (rat ChIP -seq) was scored using NRLB Models for ARNT (red), ARNT2 (blue) or NPAS4 (green), binned by motif affinity (1-10; low to high) and compared to average NPAS4 ChIP peak score (error bars represent ± standard error of the mean). **E.)** linear regression of binned affinity scores (1-10; low to high) vs mean ((±SEM) mouse NPAS4 ChIP peak scores. Linear regression p-value p < 2 x 10^-16^. Inset – mean (±SEM) peak size subset by motif sequence CGTG -, TCGTG -, CGTG +, TCGTG +, or GTCGTGA + containing ChIP peaks. **F.)** Area under receiver operator curves (±SD) for the ability of NRLB DNA binding models for ARNT (top left), HIF1α/ARNT (top right; round 3, random 18mer library), HIF2α/ARNT(bottom left; round 3, random 18mer library), or HIF2α_r2/ARNT(bottom right); round 2, FixedCore 18/22mer library) to predict ChIP-peaks from from hypoxically treated HepG2 cells NPAS4, NPAS4/ARNT, or NPAS4/ARNT2 ChIP-peaks vs size matched random sequences. NRLB models were used to score ChIP-peaks or randomly selected size matched regions to compare the ability to identify true positive transcription factor binding sites. **G.)** HIF1α or HIF2α ChIP-peak DNA (HepG2) was scored using NRLB Models for ARNT (red), ARNT2 (blue) or NPAS4 (green), binned by motif affinity (1-10; low to high) and compared to average NPAS4 ChIP peak score (error bars represent ± standard error of the mean). **H.) (left panel)** Area under receiver operator curves for the ability of NRLB DNA binding models for HIFα to predict HIF1α or HIF2α ChIP-peaks from hypoxically treated MCF7 cells or randomly selected regions. **(right panel)** linear regression of binned affinity scores (1-10; low to high) vs mean ((±SEM) HIF1α or HIF2α ChIP peak scores p-value p < 2 x 10^-16^. Inset – mean (±SEM) peak size subset by motif sequence CGTG -, CGTG +, TACGTG +, or GTACGTGM + containing HIF1α or HIF2α ChIP peaks. **I.)** (left panel) Area under receiver operator curves for the ability of NRLB DNA binding models for AhR to predict AhR, ARNT or AhR/ARNT ChIP-peaks from 2,3,7,8-tetrachlorodibenzo-ρ-dioxin (TCDD) treated MCF7 cells or randomly selected regions. (Right panel) linear regression of binned affinity scores (1-10; low to high) vs mean (±SEM) AhR/ARNT ChIP peak scores p-value p < 2 x 10^-16^.

### Information for Supplementary Figure 6

#### Flanking sequences define preferential Sim1 v HIF DNA binding

Initially, we compared kmer affinities for each TF dimer on the highest affinity IUPAC consensus, confirming highly specific DNA binding for NPAS4 and AhR on their preferred consensus sites (Supplementary Fig. 6A). In addition, it also suggested that SIM1 and HIF2α can bind preferentially to their individual consensus sequences (Supplementary Fig. 6A). We also showed that SIM1/ARNT2 preferentially bound to its top predicted 22mer DNA probe as compared to the top predicted 22mer DNA probe for HIF2α/ARNT, indicating that SIM1 favours distinct flanking sequences to HIF2α/ARNT (Supplementary Fig. 6B). Therefore, we compared upstream and downstream specificity of SIM1/ARNT2 vs HIF2α/ARNT heterodimers, demonstrating that HIFα/ARNT subunits preferentially bind TG at positions f_-3_- f_-4_ upstream of the core (NNCGTG) (Fig. 2A-G **and** Supplementary Fig. 6C). We also found that SIM1/ARNT2 bound preferentially to CT/AT sequences at f_+1_-f_+2_ downstream of the core (NNCGTG), whereas HIF2α/ARNT bound preferentially to AC (Fig. 2B **and** Supplementary Fig. 6D-E).

#### Notes on DNA shape encoded specificity upstream and downstream

bHLH-PAS transcription factors have been shown to bind flexible consensus sequences ^2, 25^. This observation, in addition to the apparent complex relationship between affinity, distinct flanking nucleotide sequences and the enrichment of AT nucleotides downstream of the Core binding site, indicated that DNA shape dependent interactions may contribute to select TF binding. Given that NRLB calculations do not specifically incorporate DNA shape parameters into models we used kmer affinity tables and NRLB models to assess shape contributions to binding affinity. Comparison of nucleotide codependences upstream of the core NNN<u>NNCGTG</u> using 12mer affinity tables indicated that all heterodimer complexes displayed preferences for trinucleotides upstream of their primary core sequences NNCGTG (Fig. 4A-D). However only NPAS4/ARNT2 and SIM1/ARNT2 were able to bind distinct pentameric cores (NNN<u>NNCGTG</u>) with similar affinity (Fig. 4B and D). This flexibility in sequence binding allowed overlapping selection for AhR/ARNT and NPAS4/ARNT2 at GTGCGTG sequences and preferential binding of SIM1/ARNT2 (vs HIF2α/ARNT) to GTGCGTG (Fig. 4E and H). Importantly, while biological settings where AhR/ARNT and NPAS4/ARNT2 are active in the same cells has not been demonstrated, AhR and NPAS4 can commonly bind and activate tiPARP in response to different stimuli and share the repeated response element GTGCGTG at the AhR/NPAS4 chromatin occupied sites^23, 26^. We hypothesized that SIM1/ARNT2 and NPAS4/ARNT2 bind to a more flexible consensus by shape directed DNA interactions. Indeed, we found increased minor groove width upstream of the core was associated with stronger DNA binding (Fig. 4F-G, 3I-J). We also found that NPAS4/ARNT2 or SIM1/ARNT2 upstream MGW-affinity relationships were similar regardless of the whether T/G or A/G was upstream of the core, respectively (Fig. 4F and I).

We also found that downstream nucleotides were associated with shape parameters. In particular, SIM1/ARNT2 and HIF2α/ARNT displayed AT enriched sequences (f_+5_-f_+8_). DNA shape-affinity profiles revealed that preferred AT-rich sequences were associated with increased affinity and decreased MGW and ProT (Fig. 4I-J **and** Fig. 3A). While AhR/ARNT and NPAS4/ARNT2 displayed downstream nucleotide preferences at similar positions to HIF2α/ARNT and SIM1/ARNT2 we did not observe the same relationships between affinity and DNA shape (discussed below).

**Supplementary Figure 6.**
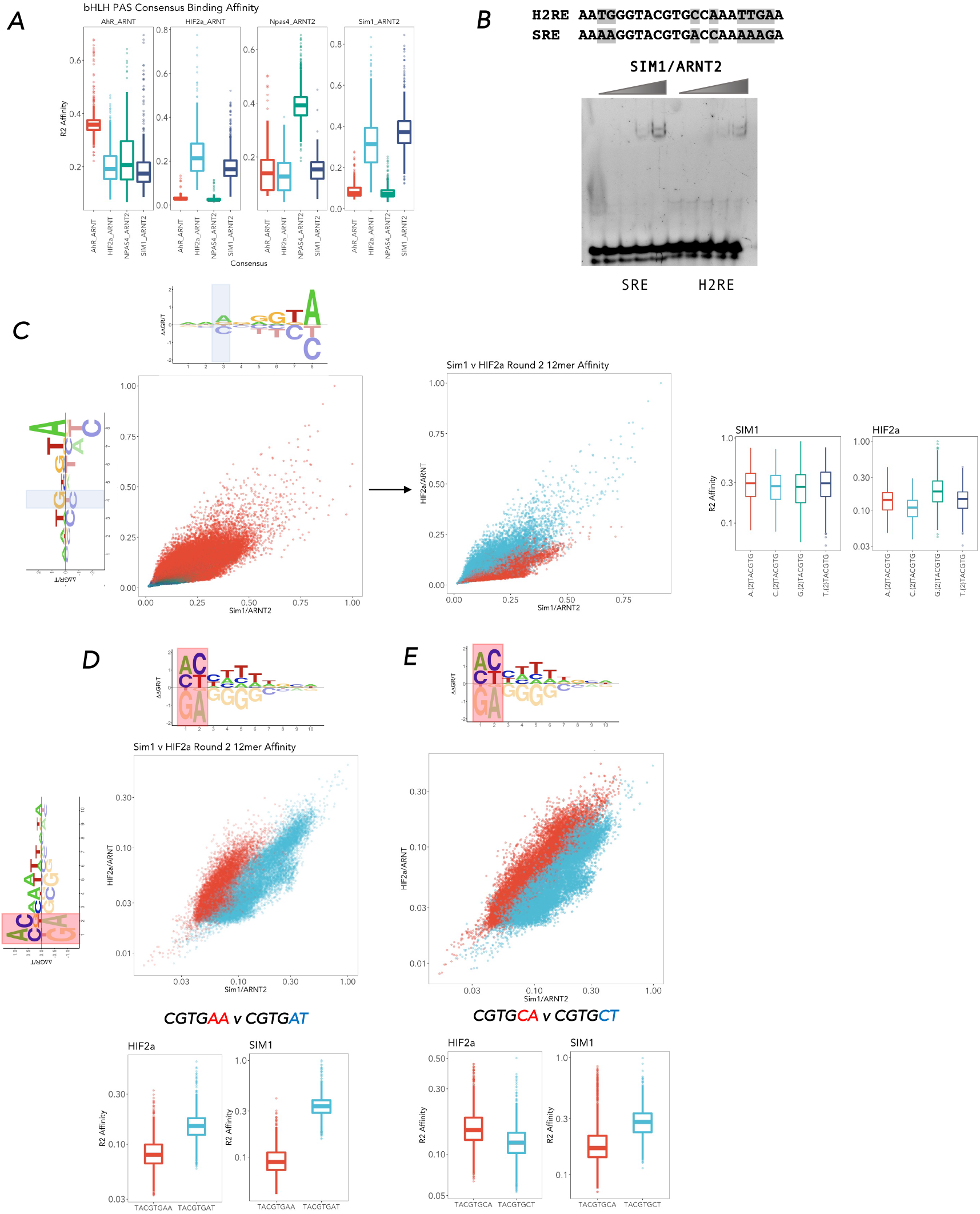
Comparison of SIM1/ARNT2 v HIF2α/ARNT DNA binding specificity encoded in Core flanking sequences. **A.)** Boxplots of Kmer affinities for the bHLH-PAS transcription factors (panels left ➔ right) AhR/ARNT, HIF2α/ARNT, NPAS4/ARNT2 and SIM1/ARNT2 from the following consensus SIM1/ARNT2 consensus = GTACGTGMY, HIF2α_ARNT consensus = GNGTACGTGM, NPAS4_ARNT2 consensus = RRDRTCGTGAY, or AhR_ARNT consensus = TTGCGTGHC (IUPAC code; R = A or G, Y = C or T, M = A or C, D = A or G or T, H = A or C or T, N = any base.) **B.)** Top Kmer sequences from modelled DNA binding sites for SIM1/ARNT2 (Sim1 response element (SRE)) or HIF2a/ARNT (HIF2a response element (H2RE)) were used in EMSA gel shift assays to confirm SIM1/ARNT2 transcription factor preference for the SRE vs H2RE. shaded nucleotides indicates variant positions **C-F.)** Transcription factor specificity encoded through the upstream flank (shaded). **C.)** Upstream nucleotide preferences shown on NRLB energy logos for HIF2α/ARNT (y) and SIM1/ARNT2 (x), scatter plot of all 12mer Kmer Affinities coloured by NCGTG (GCGTG = green, ACGTG = red) **and** scatter plot of all 12mer Kmer Affinities containing either GxxxxCGTG blue or CxxxxCGTG in red. **Right panel.** Boxplot of 12mer Kmer NxxTACGTG affinities for SIM1/ARNT2 or HIF2α/ARNT. **D-E.)** Transcription factor specificity encoded through the downstream flank (shaded). Downstream nucleotide preferences shown on NRLB energy logos for HIF2α/ARNT (y) and SIM1/ARNT2 (x), scatter plot 12mer Kmer Affinities (log_10_) selected for the presence of the indicated dinucleotide downstream. coloured by HIF2α preference (Red) or SIM1 preference (Blue). Boxplots of indicated comparisons lie below the scatterplots.

### Information for Supplementary Figure 7

#### The PAS domain encodes a novel AT-hook like domain

While structural analysis of HIF1α/ARNT/HRE and SIM1/ARNT2/HRE models indicate that a distal interaction of a PAS A loop with DNA (Fig. 2 F-G), several characteristics of the structures^11^ and SELEX-seq data led us to hypothesise that this may not be the optimal DNA-bound conformation. Firstly, nucleotide enrichment extended beyond the proposed PAS A loop-DNA contact and the likely flexible PAS A loop contains classic DNA binding residues (K and R), which may allow the loop to reorient into a more favourable (lower energy binding conformation). In addition, we also noticed that the combination of shorter DNA response elements and crystal-crystal contacts observed in structures may constrain the structural conformation of heterodimer on DNA, which may not reflect the optimal in-solution conformation in which the SELEX-seq was performed (data not shown). Given these observations we performed PAS A loop remodeling and energy minimization of bHLH-PAS heterodimer structures on longer DNA sequences derived from SELEX-seq experiments to better capture the distal DNA contacts (Fig. 4 C-E). We found that loop remodeling of HIF2a/ARNT and SIM1/ARNT PAS loops enabled the more extensive interactions with distal DNA and indicated that K and R residues within the loop come in close proximity or interact with DNA (Fig. 4C-E). We also generated structural models for NPAS4/ARNT and NPAS4/ARNT2 heterodimers on NPAS4 response elements and found that PAS A loop Arg interactions with distal DNA sites may be common to bHLH-PAS transcription factors (Supplementary Fig. 7C-E). These models appear to adopt different loop conformations which may indicate a flexible surveying of the downstream DNA and help explain why we don’t observe the same DNA Shape-affinity relationships as we do for HIF2α/ARNT and SIM1/ARNT2.

Alignment of the amino acids within the PAS loop region for Class I bHLH-PAS TF’s revealed a cluster of positively charged residues at similar positions within the loop, many of which are predicted to contact DNA. In addition, amino acids within or flanking the loop have previously been shown to disrupt DNA binding^11^ and target gene activation^27^ in AhR or HIF1α and HIF2α (DNA binding only). AT-rich binding DNA contact proteins commonly contain AT-Hook domains which interact with DNA predominantly through Arginine interactions^28^. We found that HIFα proteins display AT-hook like residues within the PAS-loop and align to known AT-hook domains within HMGA1 and MeCP2, indicating that the PAS loop may represent an unannotated AT-Hook domain (Supplementary Fig.7A-B).

#### Notes on Variants in the PAS A loop

We found several variants in clinical databases or from previous early onset obesity cohorts which are predicted to disrupt PAS-loop mediated DNA binding in NPAS4, HIF2α, or SIM1 (Supplementary Fig. 7F). In particular, we focused on Sim1.R171H variant that lay within the PAS A domain and was previously shown to be associated with early onset obesity in humans. Multiple mouse models have shown that haploinsufficiency in SIM1 results in hyperphagic obesity^7, 29–31^, however whether partial loss-of-function is sufficient to drive obesity is yet to be determined. This is of particular interest as the majority of SIM1 variants that were found to be associated with obesity resulted in modest partial loss of function (30-80% of WT)^8, 9^. As such whether these are sufficient to drive monogenic obesity or are polygenic in nature is an important unexplored question. The Sim1 variant R171H resulted in partial loss of function, as assessed by reporter assay (∼30% of WT)^9, 32^. The R171 was also shared in the paralogous gene SIM2 which also resulted in partial loss-of-function^32^. We were interested in whether SIM1.R171H was sufficient to drive monogenic obesity in humans, also addressing whether distal AT-hook interactions are important for transcription factor function. We generated a knock-in mouse model carrying the human mutation R171H by homologous recombination in ES cells, chimera generation and germline transmission. Sim1^R171H/+^ gained more weight on a high fat diet than WT mice suggesting that partial loss-of-function in SIM1 can drive human obesity.

Next, we investigated the mechanism that was underlying the SIM1.R171H loss of function. Using a SIM1 responsive reporter with the DNA binding region (bHLH-PAS A+B) of SIM1 fused to a VPR activation domain in conjunction with ARNT or ARNT2 DNA binding regions (bHLH-PAS A+B) lacking the transactivation domains we found SIM1 R171H failed to activate the reporter indicating that SIM1 R171H has a lower affinity for the reporter (Fig. 3F). In contrast SIM1-VPR and SIM1.R171H-VPR (bHLH-PAS A+B) expressed with Gal4-ARNT (bHLH-PAS A+B) activated a Gal4 responsive reporter equally (Figure 3G) indicated that loss of function is as a result of decreased DNA binding and not dimerisation with ARNT.<colcnt=7>

**Supplementary Figure 7.**
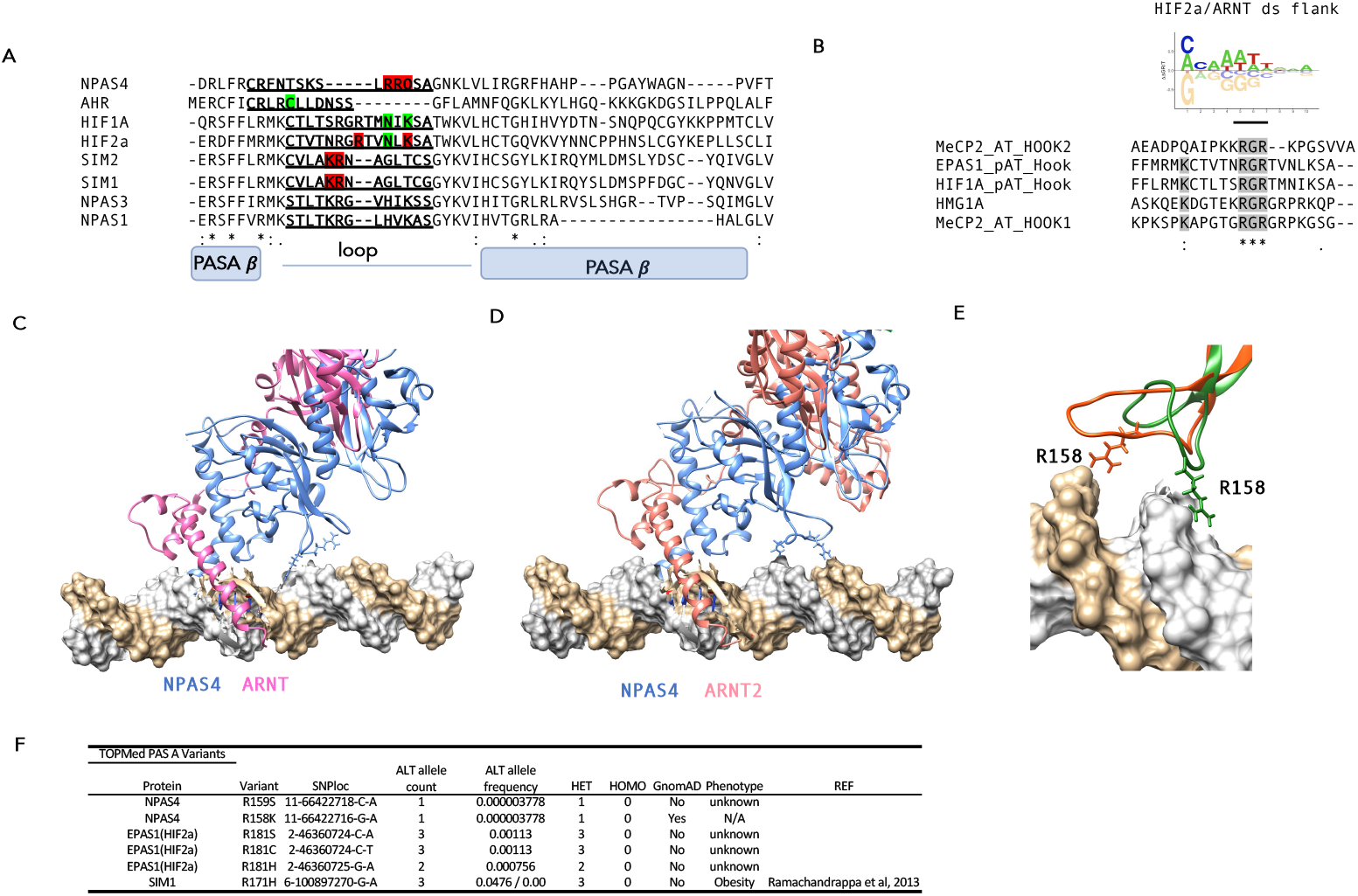
PAS A loop DNA interaction mediated by Arginine residues. **A**.) Alignments of Class I bHLH-PAS domains with basic residues in close proximity or contacting DNA from structures (or models) indicated in red and mutations leading to reduced DNA binding indicated in green. Loop regions extending from the main structure are underlined. **B**.) alignment of PAS loop from HIF1a and HIF2a with the AT-hook regions of MeCP2 (RGR motif is shaded). **C.)** and **C.)** structural models of **C.)** NPAS4/ARNT/NRE or **D**.) NPAS4/ARNT2/NRE show that the PAS loop can adopt multiple different conformation in which **E.)** Arginine residues come in close contact with AT-enriched regions of the DNA downstream of the core DNA binding site **F.)** human variants in bHLH-PAS transcription factors at DNA interacting arginine residues in the PAS-A loop. SIM1 R171H was previously identified as a variant associated with severe hyperphagic obesity.

### Information for Supplementary Figure 8

#### Comparison of wtSIM1 and SIM1.R171H interacting proteins

To further exclude confounding effects of loss of dimerisation and/or altered protein-protein interactions as an explanation for SIM1.R171H loss of function we performed affinity purification mass spectrometry to isolate SIM1 WT or SIM1 R171H interacting proteins (Supplementary Fig. 8A-E). In order to identify potential differential interactors with the SIM1.R171H region we employed three approaches. DSP crosslinking affinity purification mass spec of SIM1 WT vs SIM1.R171H using either the N-terminally truncated (Supplementary Fig. 8A) or full-length proteins (Supplementary Fig. 8B) or nuclear native (Supplementary Fig. 8C) affinity purification mass spec of full length SIM1 WT or R171H (**Supp. Table 2**). All affinity purification methods isolated SIM1 as the most abundant protein and ARNT and ARNT2 as known interacting proteins. In addition, as expected HSP90A a predominantly cytoplasmic protein was identified in whole cell extracts purifications but lowly abundant in nuclear extract purifications. We also identified many TF or chromatin binding proteins consistent with the function of SIM1 as a transcription factor.

We found that ∼97% of N-terminal interacting proteins identified as interactors of with SIM1 WT were also found in as SIM1.R171H interactors. In addition, label free quantification of interactors comparing all experiments demonstrated that SIM1 WT and SIM1.R171H displayed no difference in the ability to pull-down ARNT/ARNT2 (Fig. 3H). Quantification of SIM1 WT of R171H interacting proteins was highly consistent within each purification strategy (PCC = 0.84-0.99), and we did not identify any significantly different interactors between SIM1 WT and SIM1.R171H when comparing all interaction proteomics datasets. In addition, we confirmed that there was not differential interaction of proteins between SIM1 WT and SIM1 R171H by selecting proteins that displayed differential interaction in a single experiment and performing co-IP experiments of selected interactors (Supplementary Fig. 8E). We found that none of the interacting proteins identified by mass spec preferentially interacted with WT SIM1. Taken together we concluded that SIM1.R171H loss of function was not a result of loss of protein-protein interactions.

**Supplementary Figure 8.**
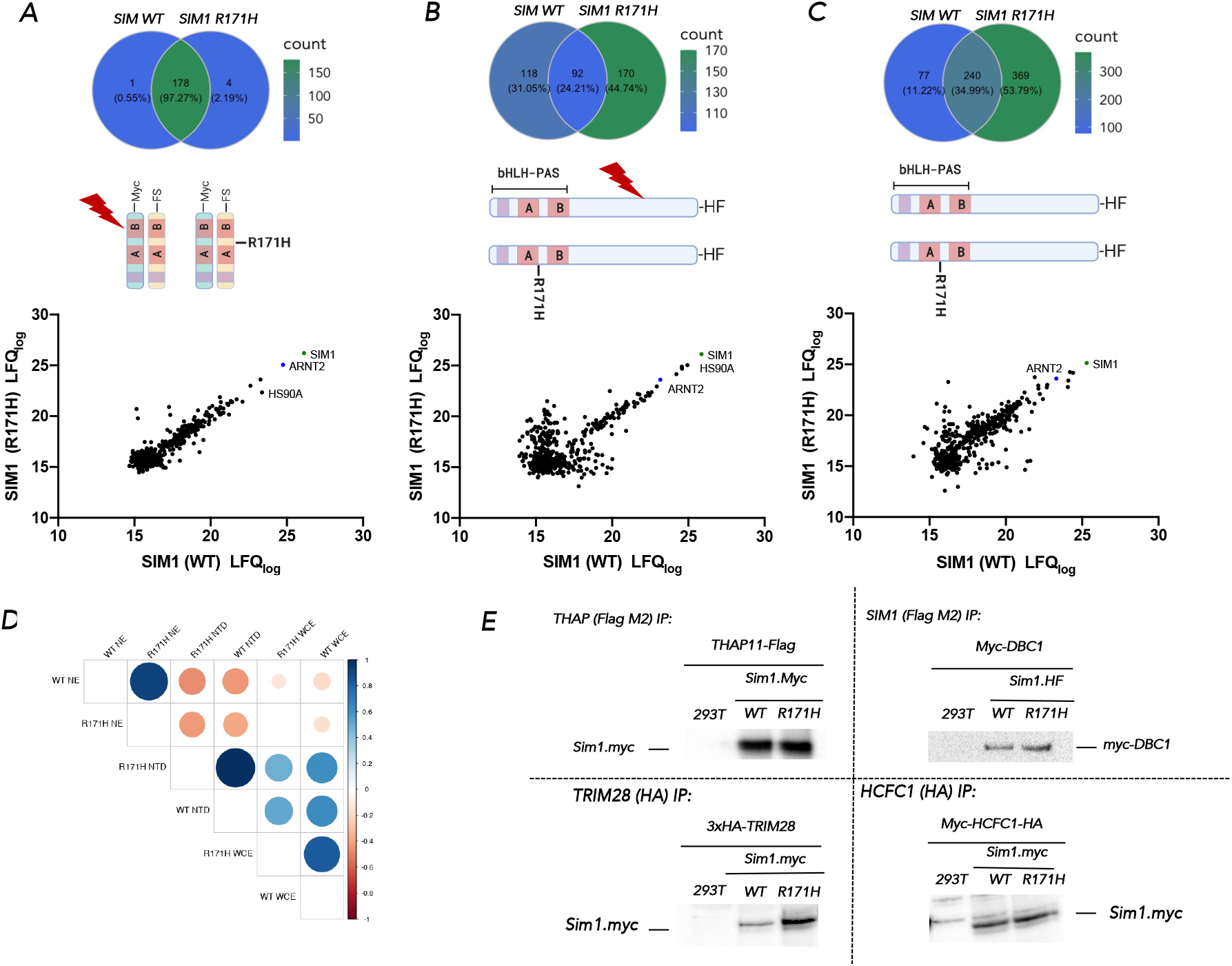
SIM1 interaction proteomics reveals similar interactome profiles for SIM1 WT vs SIM1 R171H. Three separate strategies were used to investigate SIM1 interacting proteins and identify potentially preferential interaction with WT SIM1 A.) Co-expression of N-terminal SIM1.FS and ARNT2-Myc in HEK293T cells followed by DSP in cell crosslinking and immunopurification from whole cell lysates B.) stable SIM1.HF expression in HEK293T cells DSP crosslinking and immunopurification of SIM1 complexes from whole cell lysates C.) stable SIM1.HF expression in HEK293T cells, immunopurification of SIM1 complexes from nuclear extracts. A-C.) Upper panels show Venn diagram overlap of proteins identified by mass spectrometry proteomics and schematic of strategies used to isolate complexes. Lower panel is scatter plots of Log transformed lable free quantification (LFQ) of proteins identified by mass spectrometry. D.) Correlation matrix of LFQ from each of the interaction proteomics strategies. (NE = Nuclear Extract, NTD = N-Terminal Domain, WCE = Whole Cell Extract). E.) Co-immunoprecipitation of proteins identified in interaction proteomics. Indicated proteins were coexpressed in HEK293T and Co-IP’s and western blots were performed to confirm interactions.

### Information for Supplementary Figures 9 and 10

#### DNA methylation and cell type specific DNA binding by bHLH-PAS TF

Interfamily and intrafamily DNA binding specificity and target gene selectivity appears to be mediated by sequence encoded mechanisms. However, HIF1a and HIF2a display identical DNA binding preferences, yet they have unique chromatin occupancies within the same cell types^3, 4^. HIF1a and HIF2a also display examples of cell type specificity for target gene occupancy and activation, indicating that additional mechanisms can be at play to direct transcription factor outputs. In addition, NPAS4 has been shown to have fundamental opposing roles on synapse function and gene regulation in inhibitory vs excitatory neurons^6^. We hypothesised that DNA response element methylation may play a role in directing bHLH-PAS transcription factor occupancy *in vivo*. We performed gel shifts using response elements methylated at various CpG and CpH positions within the Core response element and downstream PAS interaction site for bHLH-PAS heterodimers NPAS4/ARNT2, SIM1/ARNT2, HIF1a/ARNT, and HIF2a/ARNT (Supplementary Fig. 9A). We found that in all cases CpG and/or CpH methylation was able to reduce the affinity of bHLH-PAS transcription factors for their cognate response elements (Supplementary Fig. 9A). We found that CpG/CpH methylation of the Watson strand or the Crick strand was able to partially block DNA binding, and methylation of both strands appeared to be additive. However, we did observe that HIF1a/ARNT appeared to be more sensitive to response element methylation than HIF2a/ARNT.

To investigate whether Cp methylation directs transcription factor occupancy *in vivo*, we analysed NPAS4 ChIP seq data demonstrating a negative linear relationship between CpG methylation at ChIP peaks and NPAS4 ChIP peak intensity as well as a significantly higher NPAS4 peak score (p < 10^-9^) at unmethylated peaks vs methylated peaks (Supplementary Fig. 9B). We also found that the majority of NPAS4 sites were unmethylated in excitatory neurons but the proportion of highly methylated (>90%) sites in inhibitory neurons (parvalbumin and vasointesital peptide) was increased (Supplementary Fig. 9B). In addition, mean CpG methylation at excitatory NPAS4 ChIP peaks was significantly higher (p < 10^-16^) in inhibitory neurons compared to excitatory neurons (Supplementary Fig. 9C). Highly methylated TF binding sites were also observed at excitatory specific NPAS4 target genes (Nrp1 and Zfand5) in inhibitory neurons suggesting that DNA methylation may direct NPAS4 target gene selection (Supplementary Fig. 9D). We also found that there was an increased number of fully methylated (>90%) excitatory neuron NPAS4 target sites in inhibitory neuron cell types (Parvalbumin and Vaso-intestinal peptide neurons, Supplementary Fig. 9C). To investigate whether this was a common mechanism for cell selective chromatin occupancy for the bHLH-PAS TFs, we then compared HIF1a and HIF2a ChIP-seq data sets in MCF7, HepG2, HKC8, and RCC4 cells with DNA methylation from HepG2 or MCF7 cells. Consistently, we observed significantly higher CpG methylation at cell type specific HIFa ChIP peaks compared to shared peaks (Supplementary Fig. 10A-H). In addition, we also observed significantly lower methylation at HIF1a and HIF2a shared peaks as compared to unique peaks, but no significant difference in methylation between HIF1a and HIF2a unique peaks (Supplementary Fig. 10I-J). We found that HIF2a appear less sensitive to response element methylation than HIF1a, with the lower migrating HIF2a species less sensitive than the higher migrating species. This indicated that some select chromatin occupancy may be achieved through response element methylation. We found that while HIFa occupied sites appeared to be predominantly unmethylated, we could not identify differential methylation of HIF2a specific sites, indicating other mechanisms maybe at play. DNA methylation did appear to have a significant role in cell type specific occupancy. We observed an increased proportion of transcription factor chromatin occupied peaks which were highly methylated (>90%) in other cell types. This observation was supported by similar observations by an independent group investigating ARNT DNA binding specificity in hypoxia (as a proxy for HIFa specificity) ^33^.

**Supplementary Figure 9.**
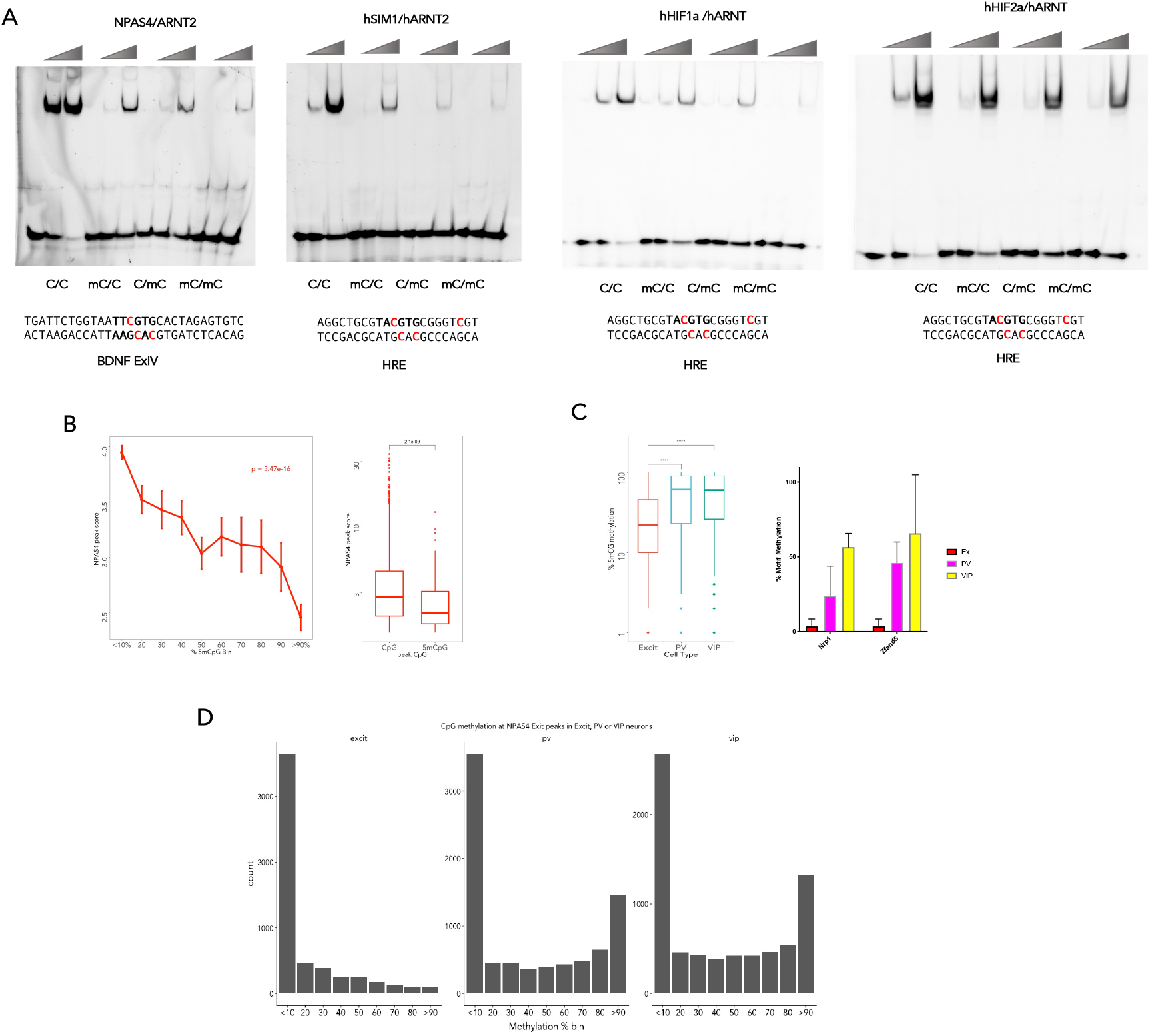
bHLH-PAS transcription factors are sensitive to response element CpG methylation, which controls cell type specific transcription factor occupancy. **A.)** NPAS4/ARNT2 on the BDNF IV promoter NPAS4 response element (**TTCGTG**), SIM1/ARNT2, HIF1α/ARNT, and HIF2α/ARNT on the hypoxia response element (**TACGTG**). Increasing amounts of purified protein was incubated with the indicated probes, which were unmethylated (**C/C**), methylated on the top strand (**mC/C**), the bottom strand (**C/mC**), or both strands (**mC/mC**) as indicated in the sequence below the EMSA gel (methylated = red). (n = 3). **B.) (left panel)** linear regression of comparing binned % CpG methylated DNA at NPAS4 occupied transcription factor sites from depolarised mouse neurons peaks to average ChIP-peak score. (model p-value is inset). **(right panel)** mean log_10_ NPAS4 ChIP peaks score at unmethylated (CpG, <10%) or methylated (mCpG, >90%) sites. **C.) (left panel)** Average CpG methylation at NPAS4 ChIP peaks (CTX, excitatory neurons, 11,344) in excitatory cortical neurons (red), inhibitory parvalbumin (PV) neurons (blue), or inhibitory vasointestinal peptide (VIP) neurons (green). Wilcoxon statistical test **** p < 10^-200^. **(right panel)** Excitatory specific NPAS4 regulated genes Nrp1 and Zfand5 are methylated at NPAS4 ChIP peaks in inhibitory neurons. Average CpG methylation (n = 2) at NPAS4 ChIP peaks (CTX, excitatory neurons) in excitatory cortical neurons (red), inhibitory parvalbumin (PV) neurons (pink), or inhibitory vasointestinal peptide (VIP) neurons (yellow). **D.)** Number NPAS4 ChIP-seq peaks (CTX, excitatory neurons) of peaks in each % CpG methylation bin (0-100%) for excitatory cortical neurons (left panel), inhibitory parvalbumin (pv) neurons (middle panel), or inhibitory vasointestinal peptide (vip) neurons (right panel).

**Supplementary Figure 10.**
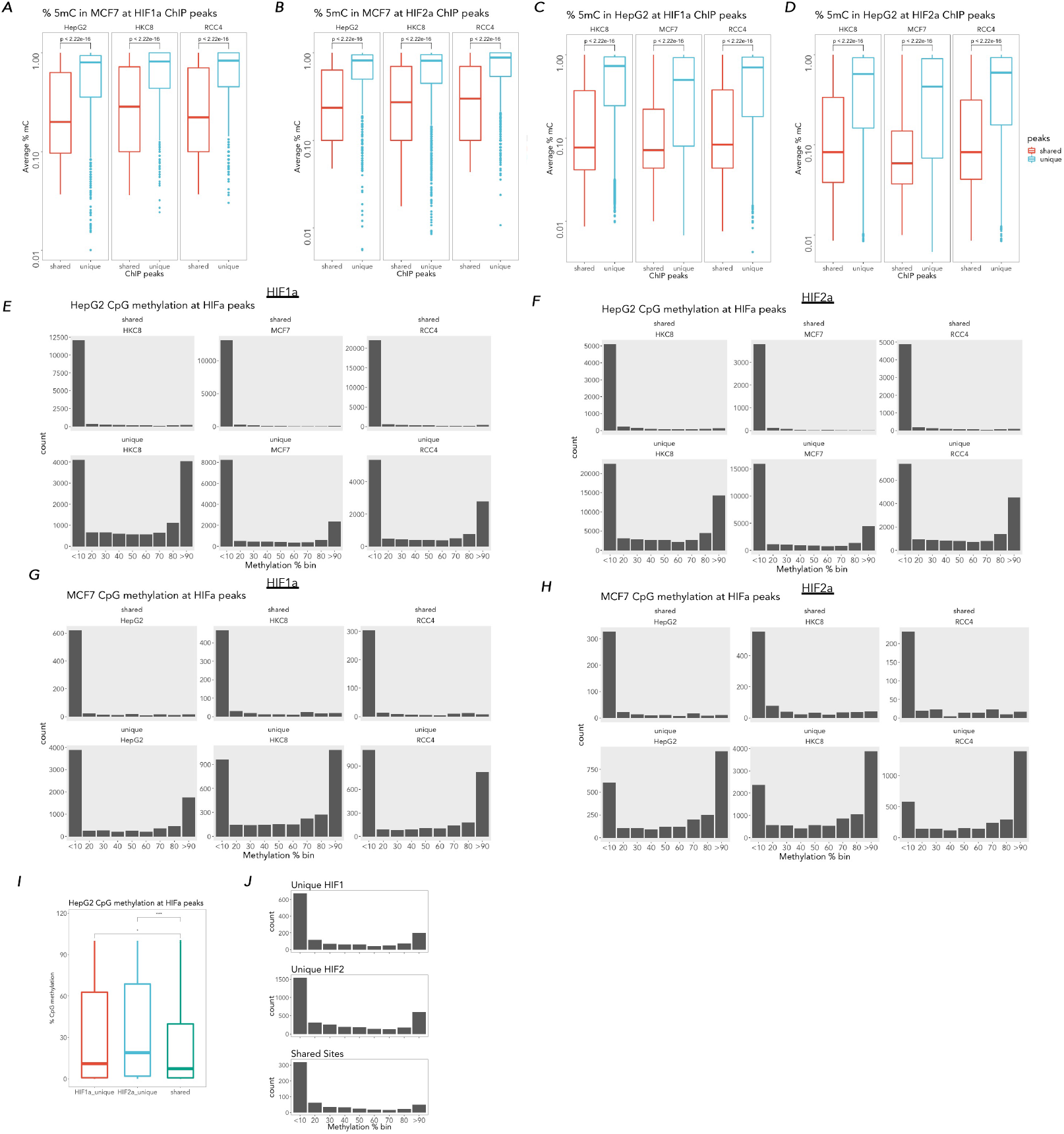
Cell type specific methylation directs bHLH-PAS transcription factor occupancy. **A.)** and **B.)** Average % CpG methylation in MCF7 cells at **A.)** HIF1α or **B.)** HIF2α unique to HepG2, HKC8, or RCC4(vhl) ChIP peaks or shared HIFα MCF7 ChIP peaks. **C.)** and **D.)** Average % CpG methylation in HepG2 cells at **C.)** HIF1α or **D.)** HIF2α unique to HKC8, MCF7 or RCC4(vhl) ChIP peaks or shared HIFα HepG2 ChIP peaks. p-values represent an unpaired comparison of %CpG methylation at common or shared peaks vs unique peaks. **E.)** The number of HIF1a ChIP peaks at HepG2 methyl CpG % CpG methylation bins (0-100%) for shared HIF1α ChIP peaks shared between HepG2 and indicated cell lines (upper panel) or unique to the indicated cell lines (lower panel; MCF7, HKC8, and RCC4(vhl)) **F.)** The number of HIF2α ChIP peaks at HepG2 methyl CpG % CpG methylation bins (0-100%) for shared HIF2α ChIP peaks between HepG2 and indicated cell lines (upper panel) or inique to the indicated cell lines (lower panel; MCF7, HKC8, and RCC4(vhl)) **G.)** The number of HIF1a ChIP peaks at MCF7 methyl CpG % CpG methylation bins (0-100%) for shared HIF1a ChIP peaks shared betweenMCF7 and indicated cell lines (upper panel) or unique to the indicated cell lines (lower panel; HKC8, HepG2, and RCC4(vhl)) **H.)** The number of HIF2a ChIP peaks at MCF7 methyl CpG % CpG methylation bins (0-100%) for shared HIF2α ChIP peaks between MCF7 and indicated cell lines (upper panel) or unique to the indicated cell lines (lower panel; HKC8, HepG2, and RCC4(vhl)). **I.)** Average percentage HepG2 CpG methylation at HepG2 HIFa ChIP peaks (HIF1α unique peaks (red), HIF2a unique peaks (blue), HIF1α/HIF2α shared peaks (green). * p-value < 1 x 10^-6^ and *** p-value < 1 x 10^-10^. **J.)** number of HIFα ChIP peaks in % CpG methylation bins (0-100%), HIF1a unique peaks (upper panel), HIF2a unique peaks (middle panel), or HIF1a/HIF2a shared peaks (lower panel).

### Final Summary

This work provides an important framework for a holistic understanding transcription factor DNA binding specificity and chromatin selectivity. Models of bHLH-PAS DNA binding generated in this work will provide improved prediction of bHLH-PAS transcription factor DNA binding sites and their affinity. In conjunction with cellular DNA methylation and chromatin features models will also allow more accurate prediction of *in vivo* chromatin occupancy and target gene identification. Importantly the bHLH-PAS transcription factor family underpins several fundamental homeostatic control pathways such as the response to low oxygen (HIF1α and HIF2α), the control of neuronal network activity (NPAS1, NPAS3, and NPAS4), the appetite regulation of body weight (SIM1), circadian timing (CLOCK and NPAS2), and signal regulated control immune (HIF1α, HIF2α and AhR) and environmental detoxification pathways (AhR). As such there are a numerous diseases, disorders and pathologies that are directly mediated by the bHLH-PAS transcription family and thus improved mapping and understanding of their DNA binding will aid in efforts to elucidate and target disease mechanisms.

In particular, this study demonstrates that a previously unknown non-canonical DNA binding of bHLH-PAS TFs though DNA shape directed mechanisms plays a major role in the sequence flexibility of SIM1/ARNT2 and NPAS4/ARNT2 transcription factors. This shape directed DNA binding also contributes to DNA binding specificity between bHLH-PAS TFs. Through investigation of DNA shape contributions to DNA binding affinity and protein DNA structures we also discovered that distal DNA contacts between a novel AT-hook domain in the PAS A loop and DNA contribute to DNA binding. Mutation of the residue predicted to make distal DNA contacts and the AT-enriched region leads to hyperphagic obesity in a mouse model outlining the importance of this distal DNA interaction to disease.

### Supplementary Methods

#### NRLB and SELEX-seq Affinity Comparisons

All comparisons of DNA binding affinities were analyzed and plotted using R. bHLH motifs were annotated using CAT-box (CATATG), CAG-Box (CAGCTG), CACC-Box (CACCTG), E-box (CACGTG), E-box-like (NNCGTG or DNCGTG (Fig 1B, Supplementary Fig 2B (ARNT2)) and random 18mer SELEX-seq affinities used to compare DNA binding specificity. In Figure 1C comparisons of SELEX-seq DNA binding affinities of MAX (round 1) vs ARNT (round 3) was on a subset of 12mer containing a palindromic E-Box sequences (fCACGTGf) and annotated by the two nucleotides upstream of the NNCACGTG flank. Figure 1D was analysed using NRLB models to score all NNNCANNTGNNN 12mers and annotated using the downstream nucleotide flanking the Core CANNTGf. Likewise, comparison of bHLH-PAS transcription factor specificity annotated and subset data to highlight differential specificity. In Fig 1C the 12mers subset for those containing sequences predicted to be most divergent in specify between SIM1/ARNT2 and HIF2a/ARNT [G|C]xxx[AGT]ACGTG[AA|AC|AT|CC|CT] and annotated by the NxxxxxCGTG (G = red and C = blue).

### Interaction Proteomics

#### Stable cell line generation

pEF-IRESpuro-hSIM1-2xHA-3xFlag or pEF-IRESpuro-hSIM1.R171H-2xHA-3xFlag were cloned and used to generate stable cell lines by transfection with 10μg of plasmid into ∼70% confluent 10cm^2^ dish of HEK293T cells, 48hrs following transfection cells were selected with 1μg/ml puromycin for 3-4 weeks until stably expressing lines were generated.

#### Whole cell full length SIM1-HF X-linking tandem immunopurification proteomics

For cross-linking mass spec we established a (dithiobis(succinimidyl propionate))(DSP) (Thermo Fisher #22585) in cell cross-linking protocol. ∼1 x 10^8^ cells per condition (parent HEK293T cells, HEK293T hSIM1-2xHA-3xFlag or HEK293T-hSIM1-2xHA-3xFlag were washed with PBS, trypsinised and resuspended in a HEPES buffered saline (HBS – 40mM HEPES pH 8.05, 150mM NaCl). A final concentration of 1mg/ml of DSP reagent was added to the cells and cross-linking proceeded at room temperature for 12mins gently rocking. Cross-linking was then quenched with a final concentration of 100mM Tris pH 8.0 and cells washed with cold PBS. Cells were then lysed with lysis buffer LB −40mM HEPES pH 8.0, 1% Triton X-100, 10mM β-Glycerophosphate, 2.5mM MgCl2, 2mM EDTA, 1mM PMSF, 2 x EDTA-free protease inhibitor pellets (Roche) at 4^0^C for 30mins. Igepal and NaCl was then added to lysates at a final concentration of 0.85% igepal and 150mM NaCl, and subsequently clarified by centrifugation at 14,000rpm for 20mins at 4°C. The resultant cross-linked supernatant was then purified by Flag M2 resin washing with 2 x lysis buffer LB, 2 x IP wash buffer (20mM HEPES pH 8.0, 250mM NaCl, 1mM EDTA, 0.1% Igepal, 1mM PMSF, 1 x protease inhibitor cocktail) and 1 x TBS (25mM Tris-HCl pH7.4, 150mM NaCl). The protein was then eluted of the resin with 3xFlag (250 μg/ml) peptide in 1 x TBS. The eluate was then incubated with HA resin (HA resin Sigma -E6779) in IP buffer, the resin was then washed 3 x with IP was buffer and eluted with 3 x 100 μl glycine pH 2 before equilibration of pH by the addition of 30μl of 1M Tris-HCl pH 9.5.

### Native nuclear tandem immunopurification proteomics

2×10^8^ cells of cells stably expressing SIM1 were used per condition for immunopurification. Briefly, cells were isolated by TEN buffer (40mM Tris pH 8.0, 10mM EDTA, 150mM NaCl), washed with PBS and cytosolic fraction isolated by resuspension of the cell pellet in 2.5x the cell pellet volume (1-2mls) of hypotonic buffer (10mM Hepes pH 8.0, 1.5mM MgCl_2_, 10mM KCl, 0.4% Igepal, 10% ficoll, 1x protease inhibitor cocktail, 1mM DTT), followed by incubation on ice for 5 mins and clarification of nuclei by centrifugation at 14,000 rpm 15mins at 4°C. Nuclei were then lysed in 20mM Hepes pH 8.0, 1.5mM MgCl_2_, 420mM KCl, 20% glycerol, 0.2mM EDTA, 1x protease inhibitor cocktail, 1mM DTT for 20mins on ice and clarified by centrifugation at 14,000 rpm 30mins at 4°C. The supernatant was then diluted to ∼280mM KCl and 0.025% Igepal with IP dilution buffer (20mM Hepes pH 7.9, 150mM KCl, 1mM EDTA, 0.1% Igepal, + protease inhibitors) and incubated with Flag M2 resin O/N at 4°C rocking. The resin was then washed 4 x with IP wash buffer (20mM Hepes pH 7.9, 250mM NaCl, 1mM EDTA, 0.02% Igepal, + protease inhibitors) and eluted with 250ng/μl 3xFlag peptide in IP wash buffer. The HA purification was performed as described above.

### N-terminal domain X-linking immunopurification proteomics

pEFIRESpuro-hSim1.3xFlag2xStrep(1-348), pEFIRESpuro-hSim1(R171H).3xFlag2xStrep(1-348) and pEFIRESpuro-hARNT2(1-455)-Myc were cloned by gibson isothermal assembly as a part of this work. 3x 15cm 50% confluent dishes of HEK293T cells per condition were used to co-transfect pEFIRESpuro-hSim1.3xFlag2xStrep(1-348) + pEFIRESpuro-hARNT2(1-455)-Myc + pNSEN (10μg/10μg/5μg = 25 μg per plate) with PEI (3:1). 60hrs following transfection the cells were cross-linked with DSP and prepared for flag purification as described above. However, the lysis buffer was modified to LB2 - (20mM HEPES pH 8.0, 1% Triton X-100, 420mM NaCl, 10% Glycerol, 1.5mM MgCl2, 0.2mM EDTA, 1 x EDTA-free protease inhibitor). Flag resin was incubated with clarified lysates and washed 3 x lysis buffer LB2 and 2x IP wash buffer II (20mM Hepes pH 8.05, 250mM NaCl, 0.02% Igepal, 1mM EDTA, 5 % glycerol). The cross-linked protein complexes were then eluted with IP wash buffer II supplemented with 250ng/μl 3xFlag peptide.

All immunopurifications were run on an SDS-PAGE gel reversing disulphide crosslinks (with 50mM DTT) and stained with either syproRuby (Thermo-Fisher, #S21900) or silver stained (SilverQuest, Thermo-Fisher LC6070). Proteomics was performed by FASP trypsin digest and cleanup, tryptic peptides were identified by Nano-LC−ESI-MS/MS an LTQ Orbitrap XL ETD MS instrument (Thermo-Fisher Scientific) or QTOF mass spectrometer (Bruker Daltonics). For the 3xFlag peptide eluted samples these were thoroughly washed on FASP vivacom 30KDa molecular weight cutoff filters prior to trypsinisation to remove 3xFlag peptide contamination.

### Co-immunopreciptation analysis

Co-immunoprecipitation of interacting proteins identified by interaction proteomics were performed essentially as described in ^34^ using Flag M2 (A2220) or HA7 (E6779) agarose resins (Sigma). pEF-IRESpuro-hSim1-2myc and pEF-IRESpuro-hSim1.R171H-2myc were described previously^32^, pEF1a-Ronin-FLAG-IRES-Neo was a gift from Thomas Zwaka (Addgene plasmid # 28020) ^35^, pcDNA Myc DBC1 was a gift from Osamu Hiraike (Addgene plasmid # 35096)^36^, pKH3-TRIM28 was a gift from Fanxiu Zhu (Addgene plasmid # 45569)^37^ pCGN-HCF-1 fl was a gift from Winship Herr (Addgene plasmid # 53309)^38^.

#### MethylC-seq Analysis

Raw FASTQ data was extracted from public repositories and 5’ read trimmed (6bp) using fastx toolkit^39^. Trimmed data was then mapped through bismark^40^ using a bisulfite-converted 1000 genomes GRCh37 (HumanG1Kv37) reference genome. Duplicates we removed using samtools markdup^41^ scheme. PileOMeth (now called MethylDackel; https://github.com/bgruening/PileOMeth) was then used to identify methylation strand bias and call CpG methylation sites based on >10x coverage per site. MCF7 datasets (3 reps (2 technical replicates) each) are available from https://www.ncbi.nlm.nih.gov//sra/?term=SRP033283 : i. SRR1036970 GSM1274126, ii. SRR1036971 GSM1274127, iii. SRR1036972 GSM1274128, iv. SRR1036973 GSM1274129, v. SRR1036974 GSM1274130, vi. SRR1036975 GSM127413. HepG2 datasets ( 2 replicates) are available from ENCODE GSE127318^42^. Mouse cortical, parvalbumin and vasointesinal peptide neuron data sets (2 replicates) from GSE63137 ^43^. Bedtools^44^ was used to map methylC sites with ChIP-peaks and intersection of peaks performed using bedtools or Granges^45^ in R.

## Notes

### Competing Interest Statement

The authors have declared no competing interest.

